# Mega-fire in Redwood Tanoak Forest Reduces Bacterial and Fungal Richness and Selects for Pyrophilous Taxa and Traits that are Phylogenetically Conserved

**DOI:** 10.1101/2021.06.30.450634

**Authors:** Dylan J. Enright, Kerri M. Frangioso, Kazuo Isobe, David M. Rizzo, Sydney I. Glassman

## Abstract

Mega-fires of unprecedented size, intensity, and socio-economic impacts have surged globally due to climate change, fire suppression, and development. Soil microbiomes are critical for post-fire plant regeneration and nutrient cycling, yet how mega-fires impact the soil microbiome remains unclear. We had a serendipitous opportunity to obtain pre- and post-fire soils from the same sampling locations because the 2016 Soberanes Fire, a mega-fire burning >500 Km^2^, burned with high severity throughout several of our established redwood-tanoak plots. This makes our study the first to examine microbial fire response in redwood-tanoak forests. We re-sampled soils immediately post-fire from two burned plots and one unburned plot to elucidate the effect of mega-fire on soil microbiomes. We used Illumina MiSeq sequencing of 16S and ITS1 to determine that both bacterial and fungal richness were reduced by 38-70% in burned plots, with richness unchanged in the unburned plot. Fire altered composition by 27% for bacteria and 24% for fungi, whereas the unburned plots experienced no change in fungal and negligible change in bacterial composition. We observed several pyrophilous taxa previously observed in Pinaceae forests, indicating that these microbes are likely general fire-responders across forest types. Further, the pyrophilous taxa that positively responded to fire were phylogenetically conserved, suggesting shared evolutionary traits. For bacteria, fire selected for increased Firmicutes and Actinobacteria. For fungi, fire selected for the Ascomycota classes Pezizomycetes and Eurotiomycetes and for a Basidiomycota class of heat-resistant Geminibasidiomycete yeasts. We hypothesize that microbes share analogous fire response to plants and propose a trait-based conceptual model of microbial response to fire that builds from Grime’s Competitor-Stress tolerator-Ruderal framework (C-S-R) and its recent applications to microbes. Using this framework and established literature on several microbial species, we hypothesize some generalizable principals to predict which microbial taxa will respond to fire.

## Introduction

The rise of the mega-fire is an anthropogenic phenomenon with unknown consequences for soil microbes and ecosystem processes. In the past, most wildfires were low severity, high frequency events (Archibald et al., 2013). These low-intensity naturally occurring wildfires helped clear away dead brush, revitalize the soil with nutrient deposition from pyrolyzed material, and assist some plants (such as serotinous pines) with their reproductive cycles (He & Lamont, 2018; Keeley et al., 2011). However, rising global temperatures (Westerling et al., 2006), elongated droughts (Mukherjee et al., 2018), and development at the wildland-urban interface (Spyratos et al., 2007), have led to the rise of catastrophic mega-fires of unprecedented size, intensity, and socio-economic impacts (Stephens et al., 2014). Whereas many Mediterranean ecosystems and plants are adapted to fire and may indeed require fire to reproduce (Keeley et al., 2011), whether plants or their associated soil microbiomes will survive mega-fires remains unknown.

The soil microbiome is an important driver of plant diversity and productivity (Van der Heijden et al., 2008), and soil biogeochemical cycling (Crowther et al., 2019). If soil microbiomes do not survive mega-fires, then associated plants, especially those reliant on symbiotic mycorrhizal fungi, may not regenerate (Collier & Bidartondo, 2009). Ectomycorrhizal fungi (EMF) and arbuscular mycorrhizal fungi (AMF) are key partners with plant roots that increase access to soil nutrients in exchange for photosynthetically derived carbon (Brundrett & Tedersoo, 2018). Moreover, soil bacteria and fungi are primarily responsible for litter decomposition (Glassman et al., 2018), nutrient cycling (Crowther et al., 2019), and soil aggregation (Lehmann et al., 2017), which are essential for ecosystem regeneration. As climate change continues to influence fire regimes with unknown consequences for ecosystem processes (Rogers et al., 2011), it is essential to determine how soil microbes that drive these processes are affected by mega-fires.

Previous studies have shown that while fire significantly decreases bacterial and fungal biomass (Dooley & Treseder, 2012) and richness (Pressler et al., 2019), fire is not totally sterilizing, and similar to plants (Keeley et al., 2011), some taxa respond positively and are adapted to fire. Fires are known to shift fungal dominance from Basidiomycota to Ascomycota (Cairney & Bastias, 2007; Pérez-Valera et al., 2018; Semenova-Nelsen et al., 2019), and many Ascomycetes are known to fruit in abundance after fire. Indeed, century-old studies describe “pyrophilous”, or fire loving, fungi from mushroom surveys (Seaver, 1909). Many of these pyrophilous fungi that fruit after fires are in the Ascomycete family Pyronemataceae (El-Abyad & Webster, 1968; McMullan-Fisher et al., 2011; Petersen, 1970) and heat-treated soil has recovered Ascomycetes in the genera *Aspergillus* and *Penicillium* (McGee et al., 2006; Warcup & Baker, 1963). Further, the Ascomycete *Neurospora crassa* is known to have heat activated spores (Emerson, 1948). While less common than Ascomycetes, Basidiomycetes can also respond positively to fire. For example, the Basidiomycete mushroom *Pholiota highlandensis* commonly fruits after wildfires (Raudabaugh et al., 2020), and the EMF species *Rhizopogon olivaceotinctus* significantly increased in frequency after a pine forest mega-fire (Glassman et al., 2016). However, less is known about the mycelial response of fungi to fire, and even less is known about bacteria, with less than 3% of studies examining microbial response to fire addressing their composition (Pressler et al., 2019). Yet, recent evidence suggests that pyrophilous bacteria may also exist (Whitman et al., 2019; Woolet & Whitman, 2020). For example, several taxa of bacteria significantly increased in frequency after a boreal wildfire, in particular the Actinobacteria *Arthrobacter* and the Proteobacteria *Massilia* (Whitman et al., 2019). Currently, most research on pyrophilous microbes is observational, and as such while we know that these taxa respond positively to fire, we do not yet know why.

While their small size and immense diversity has limited our understanding of fire effects on microbes, trait-based approaches are providing promising avenues for synthesizing microbes into suites of traits under which selective trade-offs might occur (Lustenhouwer et al., 2020; Malik et al., 2020). Trait strategies and trait syndromes are well established in the plant world including traits for wildfire response (e.g. thick bark, serotinous cones, vegetative re-sprouting) (Keeley et al., 2011; Pausas et al., 2004). Functional traits synthesize the wide diversity of species into trait groups (e.g., seed size and specific leaf area) each of which is associated with specific strategies such as longevity or drought tolerance. Due to energetic costs or evolutionary constraints associated with different functions, it is unlikely that one organism will succeed at all strategies and, therefore, will *trade off* the ability to perform one function for a less costly alternative (Reich, 2014). Identifying traits of pyrophilous microbes and trade-offs among them can improve our ability to model biogeochemical consequences such as carbon cycling under anthropogenically induced changes to fire regimes (Malik et al., 2020). A recent study hypothesized what some broad microbial fire response traits might be (fast growers, thermotolerant structure producers, and resource acquisition of post-fire nutrients) (Whitman et al., 2019). Expanding on this framework and identifying microbes that fit into these trait syndromes in a variety of biomes will provide key insights into microbial fire ecology and improve prediction of fire biogeochemical impacts. The first step in this expansion of the pyrophilous trait framework is to test the phylogenetic conservation of microbial fire response, meaning testing if more closely related organisms tend to response to fire in the same way (Martiny et al., 2013). By comparing a clade’s fire response to observations made in other fire events and in multiple ecosystems we can better predict how individual taxa will respond even if they are observed post-fire for the first time. This is because traits governing microbial phenotypic responses are often conserved within clades (Martiny et al., 2013). Thus, as the discovery of traits governing pyrophilous microbes’ fire response are uncovered, the ecological role of related microbes will be better predicted in future post-fire sampling. For example, bacterial response to both nitrogen addition (Isobe et al., 2019) and simulated global climate change factors was phylogenetically conserved across all perturbations (Isobe et al., 2020), which improves prediction of disturbance response of closely related taxa.

Most fire studies use burned areas compared to an unburned control as a substitute for not having pre-fire data from the burned region (Brown et al., 2019; Buscardo et al., 2010; Whitman et al., 2019), laboratory heating experiments (Bruns et al., 2020; Riah-Anglet et al., 2015), or prescribed fire to examine impacts on soil microbes (Brown et al., 2013; Fujimura et al., 2005). While these all provide useful information on microbial response to fire, none of these conditions replicate the size and severity of a mega-fire. Since it is unlikely to have microbial documentation before fires occur, the study of mega-fire impacts must be opportunistic. Two such studies exist, in which pre- and post-fire soil samples from the exact same sampling locations after a stand-replacing fire were available, and both were in Pine forests, and both focused solely on EMF (Baar et al., 1999; Glassman et al., 2016). Existing research on pyrophilous taxa also focuses largely on pine forests (Dove & Hart, 2017; Pressler et al., 2019), so it is unclear whether the same pyrophilous taxa will respond in other forest types or dryland ecosystems. Expanding our knowledge to non-Pinaceae forests will allow us to determine if pyrophilous taxa and their traits are generalizable or ecosystem specific.

The California redwood (*Sequoia sempervirens*) and tanoak (*Notholithocarpus densiflorus*) forest is a charismatic forest of coastal California and southern Oregon that is adapted to high fire frequency with traits including thick bark and vegetative re-sprouting (Paul Zinke, 1988; Stephens et al., 2007). Redwood-tanoak forests are also culturally significant with tanoak acorns as a staple of the Native American diet (Meyers et al., 2006). These forests are now highly threatened by both the invasive pathogen *Phytopthora ramorum*, causative agent of Sudden Oak Death (SOD), and changing fire regimes (Metz et al., 2013; Simler et al., 2018). It is possible that their associated soil microbiomes, including the AMF associated with redwoods (Afek et al., 1994) and EMF associated with tanoaks (Bergemann & Garbelotto, 2006) may help them survive these unprecedented disturbances. Yet, their wildfire response remains completely uncharacterized.

Here, we take advantage of the 2016 Soberanes Fire, a mega-fire burning >500 Km^2^, which burned through our study plots established to examine the impacts of SOD on redwood-tanoak forest microbiomes (Meentemeyer et al., 2008) (Figure 1). We were able to sample soils from two burned and one unburned plot immediately post-fire so we could assess what microbes survived the mega-fire before rain could disperse in new microbes. Because we had an unburned plot, we were able to capitalize on a before-after control impact (BACI) (Conquest, 2000) experimental design for increased inference. We thus tested the following hypotheses: H1) the mega-fire would significantly decrease bacterial, total fungal, and mycorrhizal richness in comparison to the control plot, H2) fungal composition would shift from Basidiomycete to Ascomycete dominated, and H3) pyrophilous bacterial and fungal taxa would emerge and might have similarities to previously described pyrophilous taxa from pine-forests and would fit into a trait-based conceptual model based on post-fire resource acquisition, thermotolerance, or fast growth or colonization (Whitman et al 2019). Finally, we hypothesized that H4) adaptation to fire might be a phylogenetically conserved trait, as has been indicated for microbial response to nitrogen disturbance (Isobe et al., 2019).

**Figure 1:**
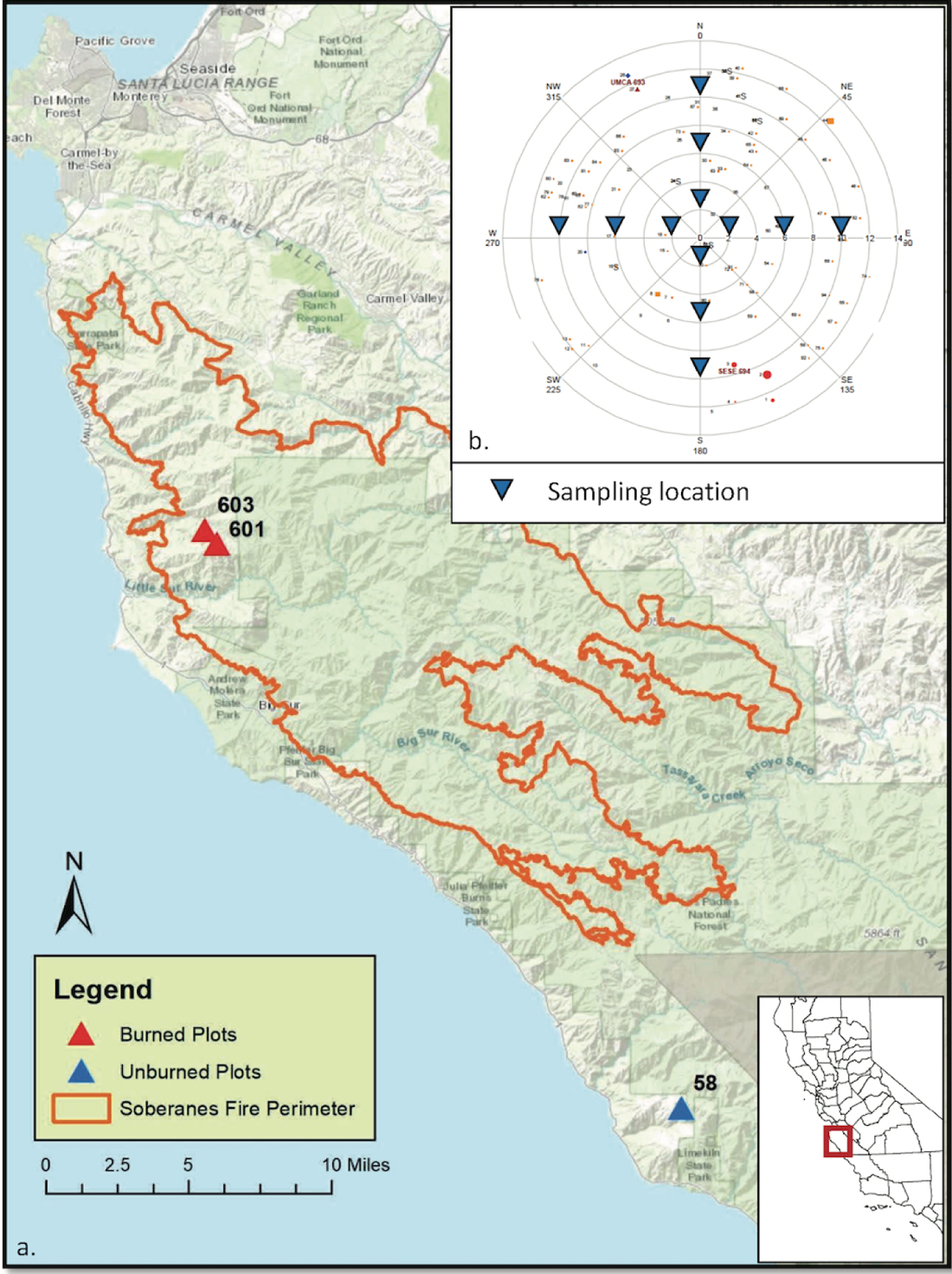
A) Map of the Soberanes Fire of 2016 with fire perimeter in orange. Triangles represent locations of individual plots in the plot network that were sampled in 2013 with burned plots in red and unburned plot in blue. B) Sampling scheme used in all plots with soils collected at 2, 6, and 10 meters from the plot center in each cardinal direction (indicated by inverted blue triangles on sampling scheme).

## Materials and Methods

### Plot Description and soil sampling scheme

In 2006 and 2007, one hundred fourteen 500 m^2^ circular plots were established in redwood forests across the California Big Sur region to study the effects of the emerging SOD outbreak (Meentemeyer et al., 2008). Big Sur has an average rainfall of 911mm/year, with most rain falling from October through April, and an average temperature range of 10.1°C to 16.1°C (Potter, 2016). On January 14^th^ and February 4^th^ 2013, soil was collected to determine soil microbial composition from a subset of those plots selected for dominance of tanoak as the only EMF host, accessibility from the road so as to keep soils on ice immediately after sampling, and similarity in slope, aspect, and elevation. We collected 12 soil cores at 2,6, and 10m from the plot center in 4 cardinal directions in an attempt to adequately sample the soil microbial communities across the entire plot (Figure 1B). The top 10cm of soil was collected using field sterilized (with 70% EtOH) Bond 8050 Releasable Bulb Planters (∼250 mL of soil). From July 22-October 12, 2016, the Soberanes Fire burned 534 km^2^ in Big Sur, burning with high severity throughout a significant portion of the fire including two of our plots (Potter, 2016)(Figure 1A). On the day that the fire was officially declared over (October 12, 2016), we were able access and collect soil samples from the exact same sampling locations (using GIS, meter tape, and notes) in one unburned (plot 058) and two burned plots (plots 601 and 603). For each plot, all tree species identification, size, and dead versus alive status were geographically mapped in 2006, 2010 and 2013 (Figures S1-S3). All plots were in UTM zone 10 and slopes ranged from 19-34°, elevations from 491-744m, and distances to the coast from 4.1-6.4km (Table S1).

### DNA Extraction and Storage

Soils were stored on ice in the field and homogenized the next day using a 2mm sieve and sterilizing with 70% EtOH between samples. For DNA extraction, 0.25g of soil was weighed and placed into tubes from the MoBio Power Soil DNA Extraction Kit and stored at 4°C. DNA was extracted within the week, following manufacturer’s protocols and stored at −20°C until analyzed.

### PCR Amplification of rRNA

Bacteria and fungi were characterized using the 515F-806R primer pair to characterize the V4 region of the bacterial 16S rRNA gene (Caporaso et al., 2011) and the ITS1F-2 primer pair (White et al., 1990) adapted for Illumina MiSeq (Smith & Peay, 2014) to characterize the ITS1 region of the fungal internal transcribed spacer region (Schoch et al., 2012). PCR recipe included 1μL of template DNA (in some cases diluted 1:10 to overcome inhibitors), 2µL of each primer at 10μM concentration, 12.5µL of Accustart Toughmix (Quantabio, Beverly, MA, USA), and 7.5µL of PCR grade water for a total reaction volume of 25µL. Thermocycler conditions began with denaturation at 94°C for 3 min; followed by 29 amplification cycles for 16S of 45 s at 94°C, 1 min at 50°C, 90 sec at 72°C, followed by a 10-min final extension at 72°C; and for ITS: 30 s at 94°C, 30 sec at 52°C, 30 sec at 68°C, with a 10-min final extension at 68°C. A mock community (ZymoBiomics, Zymo, Irvine, CA) and negative DNA extractions and PCRs were also amplified and sequenced.

### Illumina library preparation and sequencing

PCR products were pooled and cleaned as previously established (Glassman et al., 2018), with PCR products for 16S or ITS libraries pooled based on band strength from gel electrophoresis, using either 1µL, 2µL, or 3µL of the PCR product, using 3µL for the weakest bands and 1µL for the strongest. Pooled PCR products of either 16S or ITS were cleaned with AMPure magnetic beads (Beckman Coulter Inc., Brea, California, USA), quality checked for concentration and amplicon size using the Agilent 2100 Bioanalyzer (Agilent Technologies, Santa Clara, CA, USA) at the Institute for Integrative Genome Biology (IIGB) at University of California, Riverside (UCR) then pooled at a 60:40 ITS to 16S ratio. The combined library was sequenced with an Illumina MiSeq 250bp paired-end run at the UCR IIGB.

### Bioinformatics and OTU Table Construction

Demultiplexed sequences were received from the Institute for Integrative Genome Biology (IIGB) UCR core and raw reads were analyzed for quality with FastQC, and primers and barcodes were trimmed with Cutadapt v. 1.16 (Martin, 2011). We then used UPARSE v11 (Edgar, 2013) to merge forward and reverse reads with fastq_mergepairs, quality filtered with fastq_filter with fastq_maxee of 0.5, dereplicated sequences, removed singletons, and clustered 97% OTUs following established protocols (Glassman et al., 2018). For 16S, 5.2 million (M) reads were analyzed, 4.6M (88.6%) merged under the constraints for quality control, and 4.3M reads (94.4%) passed the filtering step at an expected error rate of 0.5. OTUs were then assigned taxonomic information using a RDP classifier and the GreenGenes database (DeSantis et al., 2006) (accessed 4/25/2019) in QIIME 1.9.1 (Caporaso et al., 2010). Samples identified as mitochondria, chloroplasts, or unidentified were removed, leaving a remaining 14,282 OTUs. For ITS1, 9.2M reads were analyzed, 7.5M (81%) merged, and 5.6M (75.6%) passed the filtering step. Fungal taxonomy was assigned to 97% OTUs using the QIIME 1.9.1 BLAST protocol and the UNITE database (Kõljalg et al., 2005) (accessed 5/13/2019). Samples not identified as Kingdom Fungi were removed, resulting in 3,328 OTUs. Sequences were submitted to the National Center for Biotechnology Information Sequence Read Archive under BioProject accession number PRJNA659056.

### Data Analysis

All statistical analyses and figures were produced in R 4.0.2 (R Core Team, 2020) and all scripts are available at: https://github.com/sydneyg/SoberanesFire. In order to accurately compare richness across samples with uneven sequencing depth, samples were rarefied to an even sequencing depth per sample (10,367 for bacteria and 12,089 for fungi) using the “rrarefy” function in the Vegan R package (Oksanen, 2007). In this process negative DNA extractions and negative PCRs were removed due to low sequencing depth. Mock communities were examined and removed prior to analysis. Alpha diversity metrics were calculated with the “estimate” function (observed species, ACE and Chao1) in the BioDiversityR package (Kindt, 2019) and with the “diversity” and “exp” functions (Shannon, Simpson) in Vegan (Oksanen, 2007). ACE, Chao1 and observed Species were highly correlated for both bacteria (Figure S4 A& B) and fungi (Figure S4 B&C) so all analyses and figures are based on observed species number after rarefaction. Shannon metrics were used to detect changes in evenness and Simpson was used to detect changes in dominance. Normality was tested with a Shapiro test, then ANOVA was used to test the effect of fire versus time on bacterial and fungal richness, followed by a post-hoc Tukey HSD test. Percent reduction in species richness was computed using the average species richness in each plot before and after the fire. Species accumulation curves were also constructed using “speccacum” function in Vegan (Oksanen, 2007). Richness figures were created in ggplot2 (Wickham, Chang, et al., 2019).

We used FunGuild (Nguyen et al., 2016) to analyze the effects of fire on EMF, AMF and saprobic fungi. We applied FunGuild to the unrarefied fungal OTU table, then only included “Highly Probable” guild assignments. These OTUs were then selected from the rarefied full fungal dataset using a “semi_join” function from the tidyverse R package (Wickham, Averick, et al., 2019), resulting in a total of 463 OTUs, including 222 EMF, 25 AMF, and 76 saprobic fungi. Changes in per guild species richness were then calculated.

For bacterial and fungal community composition, we calculated dissimilarity matrices as in (Glassman et al., 2018), using the “avgdist” function in Vegan (Oksanen, 2007). OTU tables were normalized by subsampling to the lowest common sampling depth 100×, then the median of the Bray–Curtis dissimilarity matrices calculated from each of subsampled OTU tables was square root transformed. We then tested the effect of time and fire on bacterial and fungal community composition with a two-way Permutational multivariate analysis of variance (PERMANOVA) (Anderson et al., 2008) as implemented with the Vegan “adonis” function. We visualized community compositional differences with nonmetric multidimensional scaling (NMDS) with the Vegan “metaMDS” function and used the Vegan “envfit” function to determine which taxa correlated well with ordination space, correcting for multiple tests with a Benjamini-Hochberg correction.

We identified pyrophilous taxa with four approaches. First, we identified taxa that correlated well with ordination space with “envfit” as described above. Second, we calculated percent changes in sequence abundance of dominant taxa (over 1% sequence abundant) before and after fire in the burned plots and visualized the most abundant taxa summarized by genera with rank abundance curves. Third, we used indicator species analysis (ISA) to identify microbial indicators in the burned plots before and after the fire using the “multipatt” function in the Indicspecies package (Cáceres & Jansen, 2019). Finally, we ran the raw OTU tables through DESeq2 (Love et al., 2019) to identify log-fold changes in abundance of each OTU in the burned plots before and after fire. We then used the DESeq2 output to determine if the taxa that positively or negatively responded to fire were phylogenetically conserved using established protocols (Isobe et al., 2019). In brief, we created circular phylogenies using maximum likelihood trees using the RAxML pipeline (Stamatakis, 2014) for 16S and ITS1and then assigned each OTU a positive or negative response to fire based on the DESeq2 analysis. Then the tree was examined for the deepest node at which >90% of the OTUs shared the same response (positive or negative). These groups were then binned into consensus clades and the mean depth of the consensus clades was calculated with consenTRAIT (Martiny et al., 2013) as implemented with the Castor “get_trait_depth” function using 1,000 permutations. The consensus clades were then mapped onto the phylogeny and colored to visualize evolutionary relationships of taxa exhibiting positive or negative responses to fire using the Interactive Tree of Life (ITOL) (Letunic & Bork, 2007). The taxonomy of clades whose response was significantly more positive or negative than expected by chance was identified with a two-tailed exact test (Mc. Donald, 2015) against the equal distribution of positive and negative responses within each taxonomic group. Because of some of the known issues with using ITS for phylogenetic analyses an attempt to strengthen our phylogeny using GhostTree (Fouquier et al., 2016) was made.

However, due to the inherent bias towards EMF in the GhostTree backbone phylogenies, aligning our OTUs to GhostTree resulted in the loss of 61.4% of OTUs despite multiple attempts to correct for this, thus rendering the GhostTree derived phylogenies not suitable for our analyses. Therefore, we decided to move forward with ITS based analysis under the guiding principle that the trends and clustering observed in our analysis, supported by literature and our other analyses, are still meaningful despite the potential weakening of statistical conclusions due to the limitations of ITS.

## Results

### Change in Microbial Richness After Fire

After rarefaction, we found a total of 12,322 bacterial and 2,878 fungal OTUs. In the burned plots (Plots 601 and 603), we found 9,116 bacterial and 1,869 fungal OTUs pre-fire, and 6,172 bacterial and 987 fungal OTUs post-fire. Fire significantly reduced both bacterial and fungal observed species richness in the burned plots, whereas richness remained unchanged in the unburned plot (Figure 2). Results did not differ if we treated samples independently (F_1,44_ = 23.75, p < 0.001,) versus if we counted them as nested within plots (F_1,1_ = 65.73, p < 0.001,). For both bacteria and fungi, number of observed taxa decreased in plot 601 by approximately 40% (37.6% for fungi and 40.1% for bacteria). In plot 603, bacterial richness declined by 52.2% and fungal richness declined by a whopping 70%. This is in contrast to the unburned plot (058) which had equivalent richness to the burned plots 601 and 603 pre-fire but experienced no change in either bacterial or fungal richness during the three years (Figure 2; Table S2). Fire also resulted in large and significant evenness reductions (Shannon diversity index decreased by 82-83% for fungi and 65-78% for bacteria) and large increases in dominance in the burned plots (measured as the inverse of Simpson’s index (Whittaker, 1965), which increased by 71-78% for fungi and by 76-82% for bacteria) with no significant change in either index for the unburned plot (Table S2). EMF richness decreased by 68% in one burned plot (plot 603), was unchanged in the other burned plot, but increased by 26% over the 3-year time span in the unburned plot (Figure S5). AMF and saprobic richness both decreased in burned plots (saprobes: 69% in plot 601 and 76% in plot 603; AMF: 60% in plot 601 and 80% in plot 603) and both increased in the unburned plot over time (saprobes: 24%; AMF: 83%; Figure S5).

**Figure 2:**
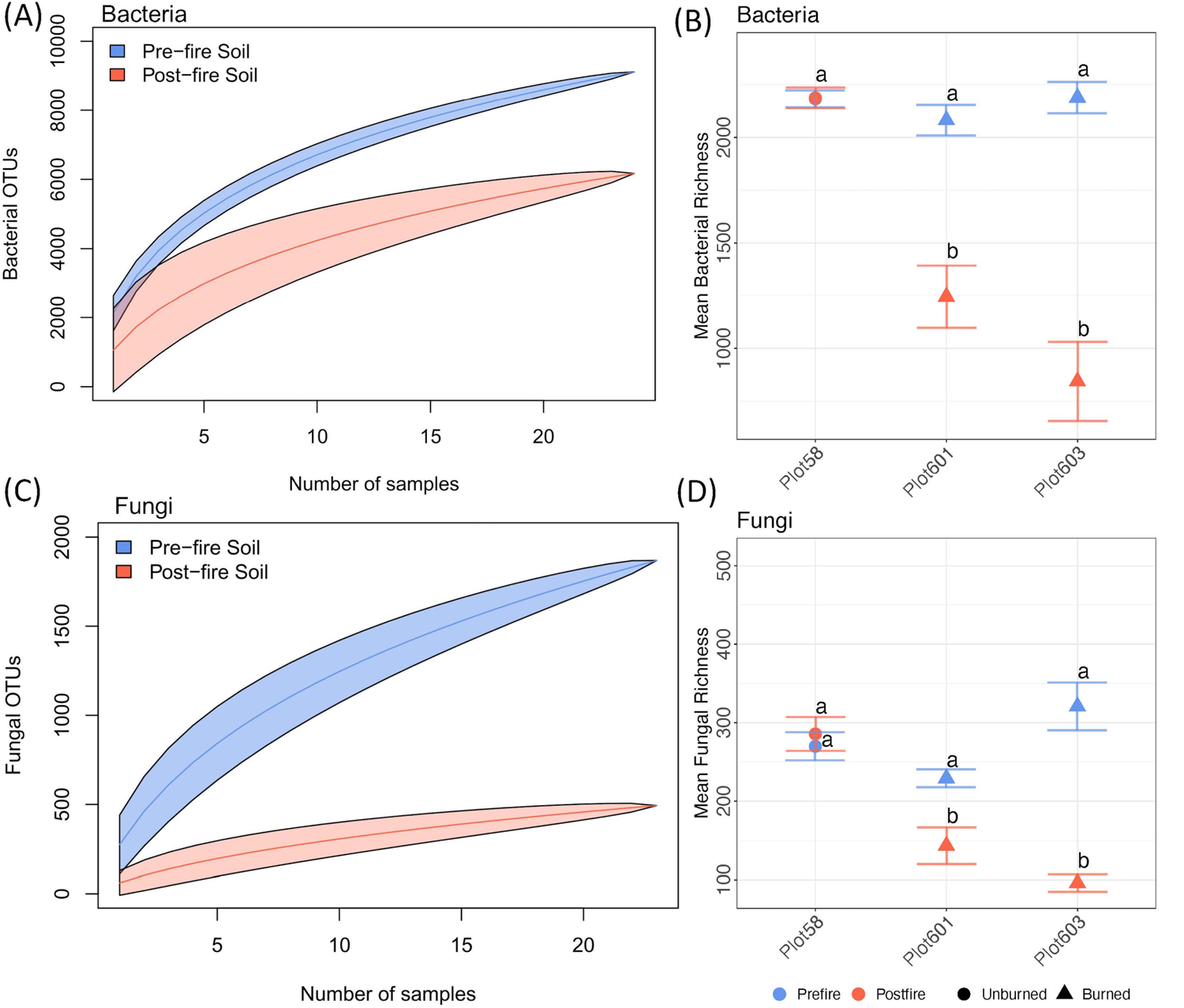
Fire reduced bacterial (A) total richness and (B) mean per sample richness in burned plots and fungal (C) total richness and (D) mean per sample richness in burned plots. Species accumulation curves represent total pre- and post-fire bacterial and fungal OTUs within the burned plots with transparency around the line representing the standard deviation. Mean per sample richness shown plus and minus the standard error. Colors differentiate sampling in 2013 pre-fire and in 2016 immediately post-fire. Shapes differentiate burned (plots 601 and 603) and unburned plots (plot 58). Statically significant difference in richness was tested using ANOVA (for burned plots, F_1,1_ = 65.73, p < 0.001, for unburned F_1,1_ = 0.005, p = 0.943). Letters represent Tukey HSD differences.

### Change in Microbial Composition After Fire

Fire resulted in a large and significant change in bacterial (Adonis R^2^ = 0.27, p < 0.001) and fungal (Adonis R^2^= 0.24, p < 0.001) community composition, while the unburned plot experienced no change in fungal composition and a small change in bacterial composition (R^2^= 0.09, p < 0.01; Figure 3, Figure S6). Compositional changes in bacteria were largely driven by increases in the Actinobacteria and Firmicutes phyla post-fire and decreases in the Proteobacteria, Gemmatimonadetes, Verrumicrobia, Chloroflexi, Elusimicrobia, Planctomycetes, Acidobacteria, Bacteroidetes, and Saccharibacteria phyla (Figure 3B). Compositional changes in fungi were driven by large increases in the Basidiomycete genus *Basidioascus* and the Ascomycete genus *Penicillium* and decreases in the Mucoromycota genus *Mortierella*, and the Ascomycete genera *Ilyonectria*, *Metarhizium*, *Cladophialphora*, *Pectenia*, *Humicola*, and *Exophiala* (Figure 3D).

**Figure 3:**
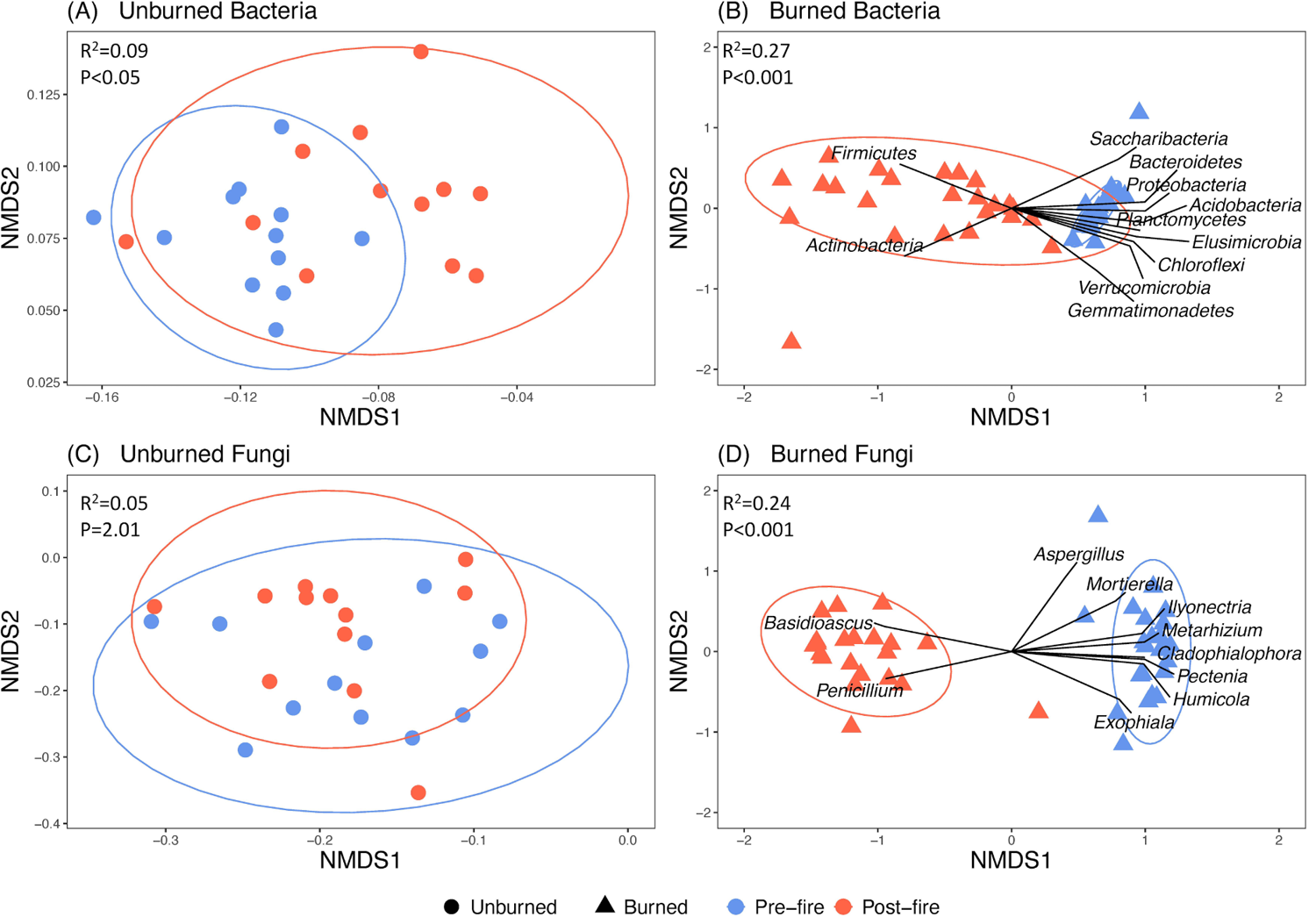
NMDS of Bray-Curtis Dissimilarity ordinations comparing bacterial composition A) in unburned plot and B) burned plots and fungal composition in C) unburned plot and D) burned plots with colors indicating the 2013 pre-fire and 2016 post-fire samplings shapes differentiating burned and unburned plots. Ellipses represent 95% confidence interval from the centroid for each group. Notice the difference in scales for all NMDS with much smaller scales for NMDS axes in unburned plots representing a much smaller degree of compositional turnover than for burned plots with much larger axes. Adonis R^2^ and p-values represent the difference in composition for pre-versus post-fire. Envfit model depicts which taxa are driving the changes in community composition for B) bacterial or D) fungal communities. The top phyla contributing to change are shown. Identification of the top contributors was done by R^2^ value after constraining p value at less than 0.001 and adjusted p-value to less than 0.01.

### Change in Relative Sequence Abundance of Bacterial Taxa After Fire

Pre-fire, bacterial OTUs dominating (≥1% of sequence abundance) the burned plots were primarily Proteobacteria (84.4%) and Acidobacteria (15.6%) with the most dominant taxa, Proteobacteria species in *Bradyrhizobium* and *Rhodoplane*s, occupying around 2.3% sequences each (Figure 4A; Table S3). Post-fire, there was a shift in the dominant phyla (≥1% total sequences), with a loss of Proteobacteria and a dominance of Firmicutes (82%) and Actinobacteria (18%). The most abundant taxon also had an increase in dominance, a Firmicute in the genus *Sporosarcina*, dominating 6.9% of the sequences, followed by Firmicutes genera *Fictibacillus, Thermoflavimicrobium, Bacillus, Solibacillus, Cohnella,* and Actinobacteria genera *Micromonospora, Pseudonocardia,* and the family Micromonosporaceae (Figure 4A).

**Figure 4:**
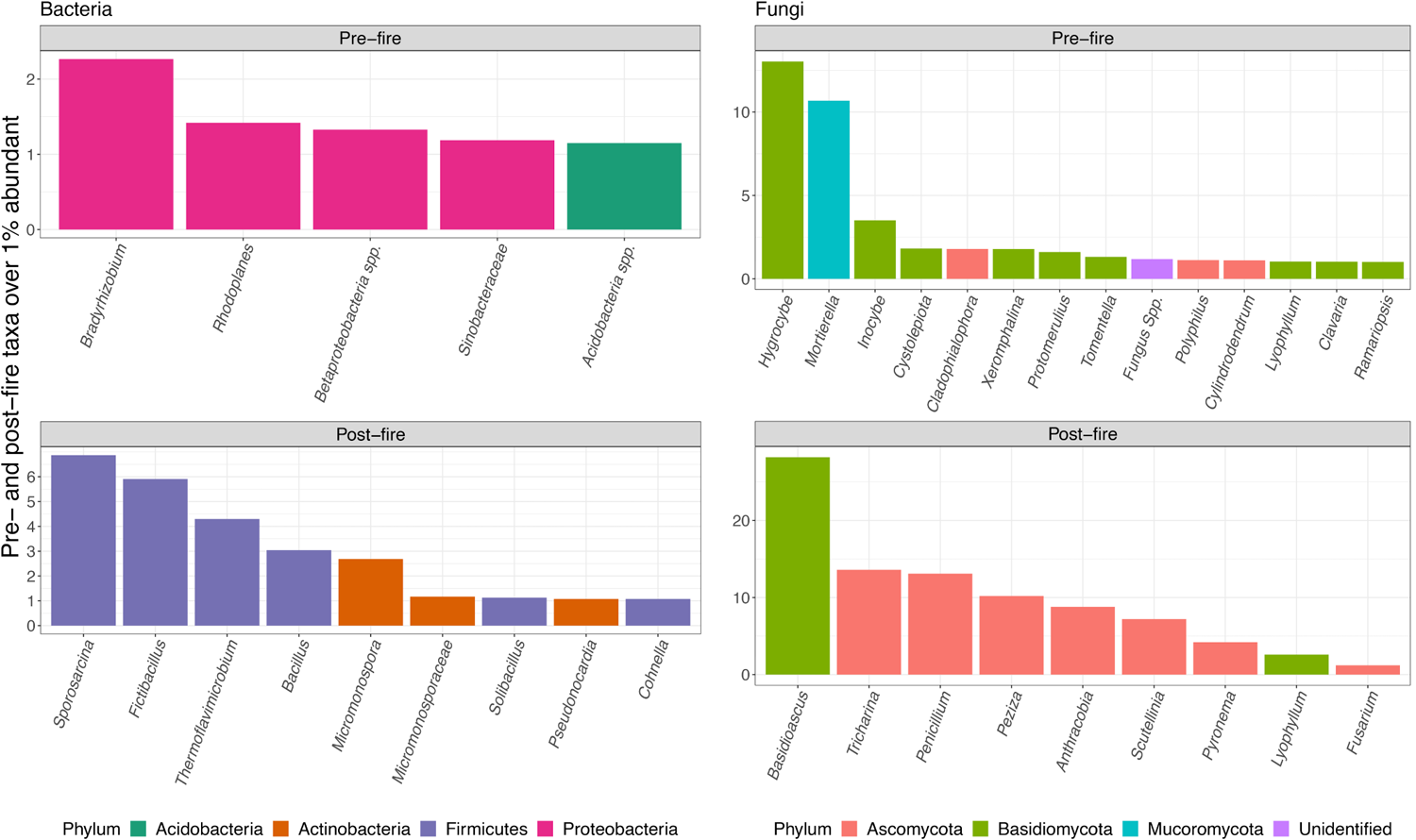
Rank abundance curves of taxa ≥1% sequence abundant for bacteria and fungi pre-fire (top) vs. post-fire (bottom) grouped by genus and colored by phyla. Where genus level identification could not be determined, a higher order of classification is given.

### Change in Relative Sequence Abundance of Fungal Taxa After Fire

Pre-fire, fungal taxa ≥1% sequence abundance were dominated 62% by the phylum Basidiomycota, followed by 25% Mucoromycota, 10% to Ascomycota, and 3% unidentifiable to phylum (Figure 4B; Table S4). Pre-fire, the most dominant fungi were the Basidiomycete *Hygrocybe acutoconica* (5.94%) and the Mucoromycete *Morteriella baineri* (4.62%). Post-fire, among the most abundant taxa, Ascomycota dominated (65%), with Basidiomycota falling to 35%, and a complete loss of the Mucoromycota. There was also a large increase in dominance with the most abundant taxon post-fire, a Basidiomycete Geminibasidiomycete yeast *Basidioascus undulatus*, accounting for 28% of the sequences (Figure 4B). The other top taxa were dominated by genera in the Ascomycota *Penicillium* 13.1% and *Fusarium* 1.2% and the family Pyronemataceae (*Tricharina* 13.6%, *Peziza* 10.2%, *Anthracobia* 7.6%, *Scutellina* 7.2%, *Pyronema* 4.2%).

### Bacterial indicator species

Bacterial indicator species analysis (ISA) revealed 86 indicator OTUs shared between the burned plots pre-fire, and 21 OTUs indicative of the post-fire community (Table S5). The top ten taxa identified for the post-fire grouping were two OTUs from the Firmicutes family Thermoactinomycetaceae, one OTU from the Actinobacteria family Thermomonosporaceae, OTUs from the Firmicutes genera *Thermoflavimicrobium, Fictibiacillus, Cohnella, Paenibacillus, Bacillus, Tepidibacterium,* and the Actinobacteria genus *Streptomyces*.

### Fungal indicators species

Fungal ISA identified 13 indicator taxa for the burned plots pre-fire and 5 indicator taxa post-fire (Table S6). The 13 pre-fire taxa identified were 4 species from the Mucoromycota genus *Mortierella* and the rest Ascomycetes belonging to the genera *Polyphilus, Pycnopeziza, Thelonectria, Pectenia, Cladophialophora,* and *Phomopsis*. The five fungal indicators of the post-fire group were all Ascomycetes, one in the Aspergillaceae, *Penicillium decumbens*, and the rest in the Pyronemataceae: *Anthracobia* sp., *Geopyxis alpina, Peziza vacinii*, and *Tricharina praecox*.

### Phylogenetic conservation of pyrophilous taxa

Lineages of both bacteria and fungi that negatively or positively respond to fire appear to be phylogenetically conserved at the class level for fungi and the phylum level for bacteria (Figures 5 & 6). There is strong clustering of bacteria positively responding to fire in the Firmicutes and Actinobacteria and a few positive responders in the Acidobacteria (Figure 5; Table S7). All other phyla belonged to consensus clades that either did not show the patterned response to fire or responded negatively. For fungi, the lineages that positively respond to fire include the Ascomycete class Pezizomycetes (containing Pyronemataceae) and the Basidiomycete class Geminibasidiomycetes (containing *Basidioascus*) (Figure 6; Table S8). There are also positive interactions sprinkled within some of the other classes, which appear to be conserved at the order level rather than class level, namely the Eurotiomycetes order Eurotiales (contains Aspergillaceae) and the Agaricomycete order Russulales. All other groups belonged to consensus clades that responded negatively or did not show the phylogenetic patterned response.

**Figure 5:**
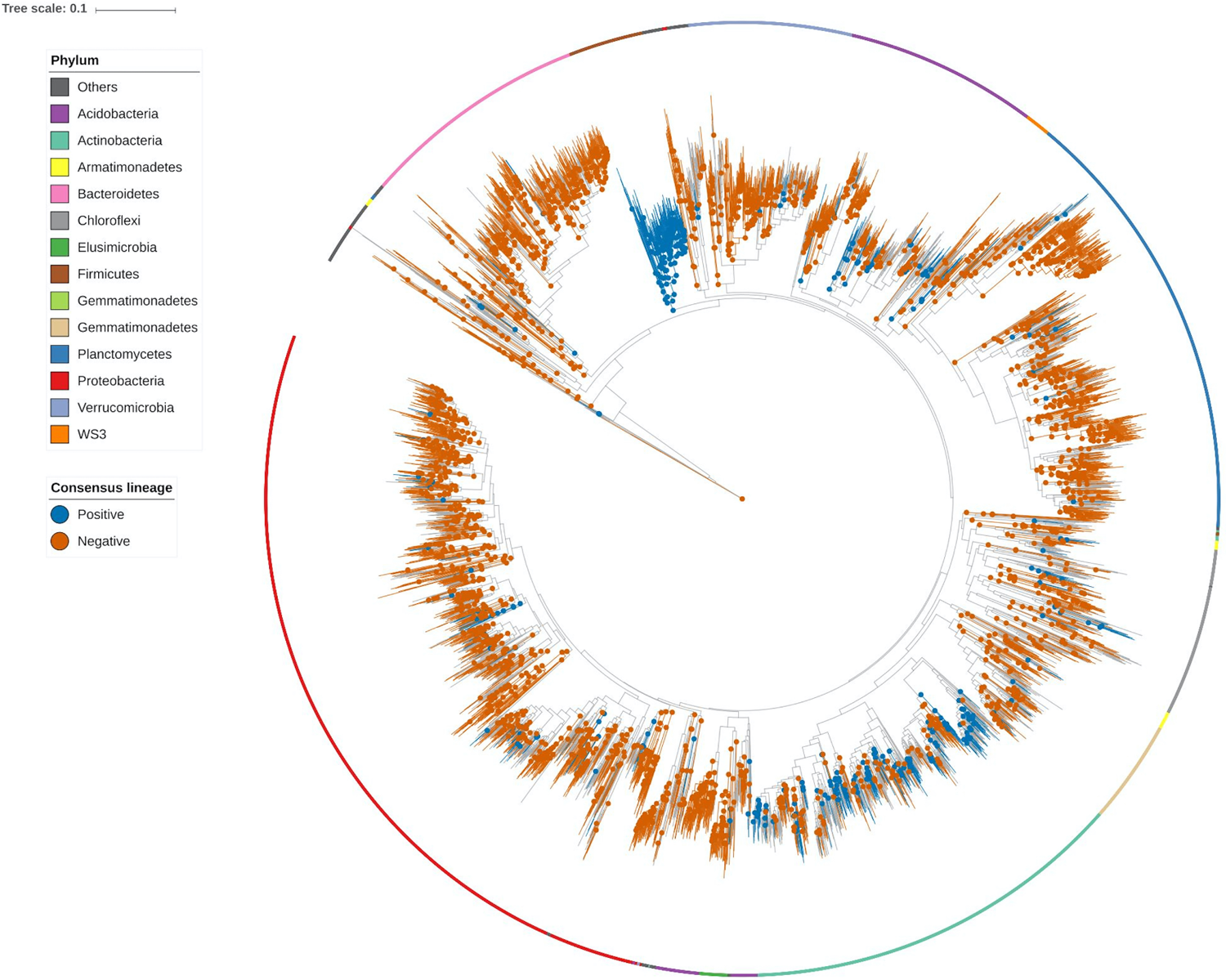
Circular phylogeny of all bacterial OTUs observed based on 16S rRNA. Consensus Lineage refers to the response to fire as measured using DeSEQ2 analysis. Lineages are colored based on positive (blue) or negative (orange) response to fire. The colored bars circling the outside of the phylogeny correspond to each OTUs respective phyla.

**Figure 6:**
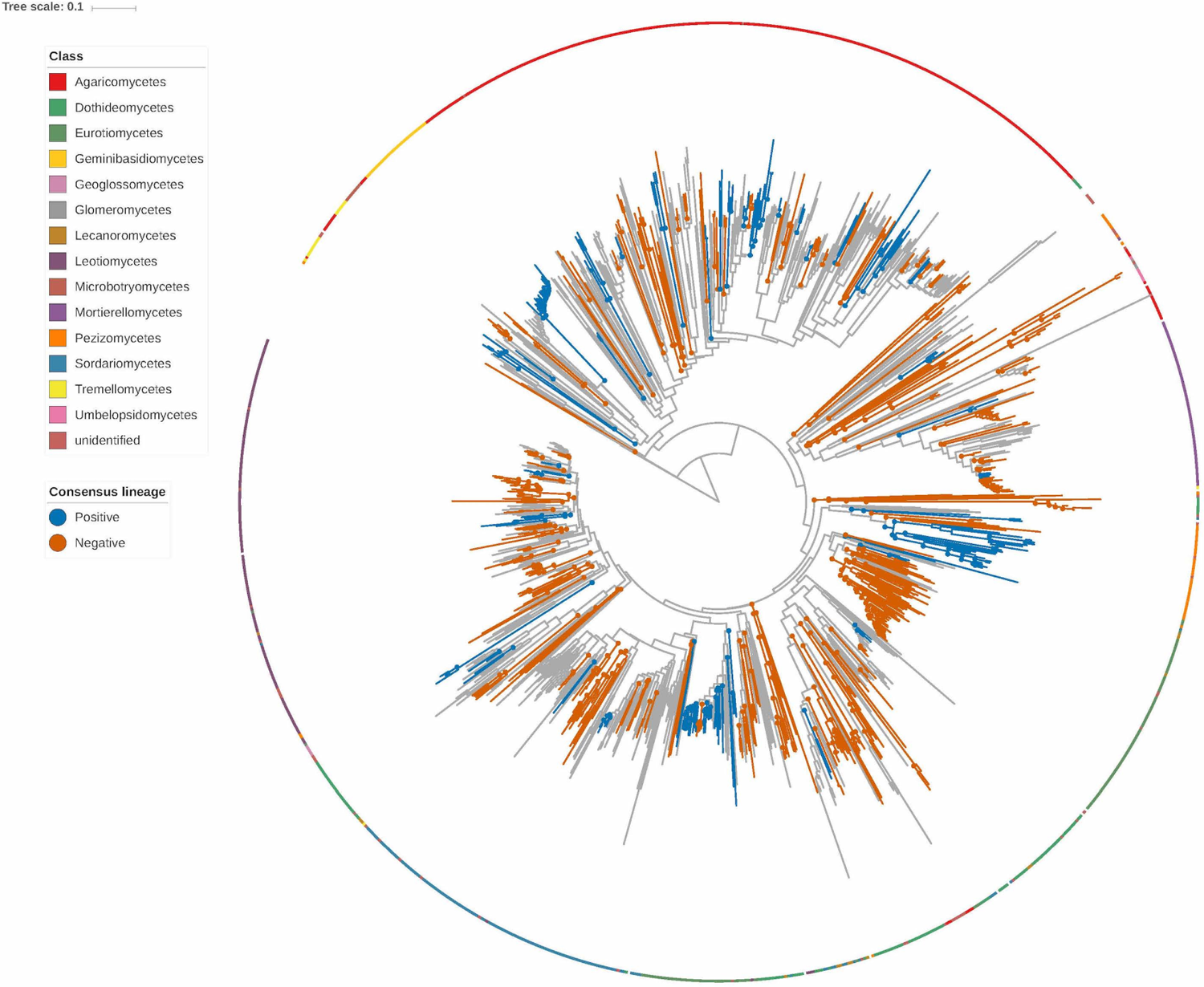
Circular phylogeny of all fungal OTUs observed based on ITS1. Consensus Lineage refers to the response to fire as measured using DeSEQ2 analysis. Lineages are colored based on positive (blue) or negative (orange) response to fire. The colored bars circling the outside of the phylogeny correspond to each OTUs respective class.

## Discussion

Here, we take advantage of the 2016 Soberanes Fire burning through our plots and show for the first time the effects of a mega-fire on bacterial and fungal communities with a rare pre- and post-fire dataset in a redwood tanoak forest. In accordance with our hypotheses, the mega-fire significantly decreased bacterial and fungal richness (H1) (Figure 2); fungal composition shifted from Basidiomycete to Ascomycete dominated (H2) (Figure 4); many pyrophilous taxa previously identified from other biomes appeared in a redwood tanoak forest and are likely generalizable and may fall into trait categories analogous to plants (H3); and adaptation to fire is likely a phylogenetically conserved trait across bacteria (Figure 5) and fungi (Figure 6) (H4).

We found that the mega-fire led to large and significant reductions in bacterial and fungal richness. Reduction in microbial richness is a typical result after fire (Brown et al., 2019; Day et al., 2019; Pérez-Valera et al., 2018; Whitman et al., 2019), however the degree to which fire affects richness varies. In our study, fire reduced the average species richness per sample by up to 52% for bacteria and up to 70% for fungi (Figure 2). In contrast, a meta-analysis of fungal response to fire found an average richness reduction of 28% (Dove & Hart, 2017). However, the range for both fungal (12-80%) (Brown et al., 2019; Pulido-Chavez et al., 2021) and bacterial (20-58%) (Brown et al., 2013; Pérez-Valera et al., 2018) richness reduction can be quite large.

This is likely due to differences in fire severity, which is often not measured or consistently reported in post-fire microbial surveys, despite that more severe fires have been shown to have greater impacts on the soil community (Whitman et al., 2019). Moreover, when reported, it is based on plant mortality measured at coarse levels (30km^2^ in the case of the Soberanes Fire) (Potter, 2016) that do not correspond well to the scale of the soil core and can lead to wide disparities in the presence or absence of live plants, duff, or ash in soil cores all taken in high severity defined fire zones. We also found that EMF richness declined following fire, which makes sense given large host mortality, and is in accordance with other high severity fires in pine forests (Glassman et al., 2016; Pulido-Chavez et al., 2021; Reazin et al., 2016). This large-scale reduction in the microbial richness can have far reaching impacts on the surrounding ecosystem from radically altered nutrient cycling (Crowther et al., 2019) to the inability to reestablish critical plant species (Van der Heijden et al., 2008).

The mega-fire led to large compositional shifts in both bacterial (27%) and fungal (24%) communities (Figure 3). Studies often report significant changes in fungal compositional turnover post-fire but variable change in bacterial composition (Pérez-Valera et al., 2018; Pressler et al., 2019). However, sampling immediately post-fire does reveal significant reductions in bacterial richness and evenness (Brown et al., 2019; Ferrenberg et al., 2013) and one study showed with increasing fire severity the reduction in richness and evenness becomes more drastic (Brown et al., 2019). Depending on the biome, changes in microbial composition can be as short lived as 21 days after simulated soil heating (Bárcenas-Moreno & Bååth, 2009) or as long as 19 years post-fire in a Spanish Mediterranean ecosystem (Pérez-Valera et al., 2018). A study in a Chinese pine forest speculated that bacterial composition would recover within one growing season based on data from 6 months post-fire (Li et al., 2019). This disparity in resilience is likely due to differences in fire severity and timing of post-fire sampling. Fires that are not as severe often produce smaller shifts in bacterial and fungal composition (Buscardo et al., 2010). The effect of fire on microbial richness and composition is also likely more transient in some biomes and longer lasting in others. As such, studies that sample 1-2 years post-fire may miss changes in bacterial and fungal communities depending on the biome. In the case of our data, these observed changes stemmed from a severe mega-fire where the changes to the landscape can be quite drastic and took place in a well-established forest and therefore recovery could take much longer.

Changes in dominance can be important indicators of how post-fire microbial communities assemble, as has been documented in plant communities (Moser & Wohlgemuth, 2006). Fires often lead to shifts in plant community evenness to increased dominance by specific fire-adapted species such as long leaf pine (Glitzenstein et al., 1995), *Ceanothus* shrubs (Lawson et al., 2010), or Manzanita in chaparral systems (Vogl & Schorr, 1972). These effects can be more pronounced when fire severity increases (Kuenzi et al., 2008). Fire similarly alters microbial community assembly processes (Ferrenberg et al., 2013). Here, we find patterns of fire induced-dominance in microbial communities, with a single fungal taxon (*Basidioascus undulates*) dominating 28% and a single bacterial taxon (*Sporosarcina spp.*) dominating 7% of the sequences post-fire. While changes in dominance have been documented in microbial communities post-fire (Pérez-Valera et al., 2018; Whitman et al., 2019), they are at lower levels (2-4% for most abundant taxon) likely because they sampled later after the fire and after the first rains post-fire thus obscuring initial compositional shifts. An experimental fire in laboratory “pyrocosm” (a small-scale highly controlled experimental fire to test the effects of fire on soil samples) similarly found huge increases in dominance with the most abundant taxon *Pyronema domesticum* occupying 57% of the sequences within 2 weeks of the fire (Bruns et al., 2020). We interpret this level of dominance to indicate the opening of niche space due to massive microbial death of the pre-fire dominants that allows the few microbial taxa that are thermotolerant or capable of capitalizing on post-fire resources or eating microbial necromass to take over.

After fire, we found a complete turnover in the abundant bacterial taxa (>1% sequence abundant) from Proteobacteria and Acidobacteria domination to Firmicutes and a few Actinobacteria (Figure 4). These changes are likely driven by thermotolerance of endospore forming Firmicutes, which have also been found to increase in abundance following laboratory heat treatments of soil samples (Filippidou et al., 2016; Jurburg et al., 2017) and soon after Mediterranean fires (Pérez-Valera et al., 2020). Post-fire environments are often characterized by increases in pH (Neary et al., 1999), which may favor both thermotolerant and alkaline tolerant Actinobacteria (Shivlata & Tulasi, 2015). Of the specific genera dominating our communities post-fire, only *Thermoflavimicrobium* (Yoon et al., 2005) and *Sporosarcina* (Jurburg et al., 2017) have previously been shown to increase after laboratory heating. Other studies have shown an increase in Actinobacteria and Firmicutes post-fire, though only the *Bacillus spp.* in our study is a potentially shared taxon with those other studies (Ferrenberg et al., 2013; Li et al., 2019; Whitman et al., 2019). In contrast, most other groups of bacteria negatively responded to fire, or only responded positively sporadically within a clade (Figure 5). Lineages can have differing responses within a clade when lineages challenged with the same evolutionary pressures do not respond the same way (Isobe et al., 2019). Furthermore, the degree to which a trait is conserved within lineages may reflect the genetic complexity of the response or trait, with more complex responses being less conserved due to needing many pathways to be conserved across all taxa of a lineage (Isobe et al., 2019). It may be then that the genetic drivers of Firmicutes’ response to fire are either less genetically complex or more genetically integral to that phylum while Actinobacteria’s response may be genetically varied or require the conservation of a greater number of genetic pathways in order to successfully adapt to fire. Finally, as this study is among the first to comprehensively characterize the soil microbiome of the charismatic redwood-tanoak forests, we note a few similarities in our pre-fire soils with a recent bacterial study of coastal redwood roots which also showed the most abundant taxa as *Bradyrhizobium* and *Rhodoplanes* (Willing et al., 2020). Additionally, a study of the microbial composition of the coastal redwood’s sister genus, the giant sequoia (*Sequoiadendron giganteum),* also showed domination by *Bradyrhizobium* and Sinobacteraceae sp. of the Proteobacteria (Carey et al., 2019). The similarities with the giant sequoia soil communities also extend to fungi, with *Hygrocybe* dominating both our redwood forests pre-fire and in the giant sequoia forest.

After the fire, the fungal communities were dominated (28%) by the yeast *Basidioascus undulates,* which was rare (0.05%) pre-fire*. Basidioascus* and *Geminibasidium* are newly described heat-resistant and xerotolerant Basidiomycete yeasts isolated from laboratory heat-treatments to soil (Nguyen et al., 2013). *Basidioascus* was originally described as an Ascomycete because their spores were spuriously described as single-spored ascii. Perhaps the visual similarity to Ascomycetes is representative of convergent evolution with other ascomycetes, which are more often associated with fire (Cairney & Bastias, 2007). Only one other wildfire study has found *Basidioascus* to increase post-fire (Pulido-Chavez et al., 2021) though not with the level of dominance in our study. A couple recent studies have found an increase in dominance of the sister genus *Geminibasidium*, including a study of a *Pinus ponderosa* forest in the American Pacific Northwest (Pulido-Chavez et al., 2021) and a study of *Larix* and *Betula* dominated forests in northeastern China (Yang et al., 2020). *Geminibasidium* was also present at 0.05% sequence abundance, the same amount as our pre-fire soil, in an unburned giant sequoia forest (Carey et al., 2019). While Geminibasidiomycetes are positively selected for by fire (Figure 6) likely due to their thermotolerance and xerotolerance (Nguyen et al., 2013), they have been missed by existing descriptions of pyrophilous fungi that are mainly based on fruiting body surveys (McMullan-Fisher et al., 2011).

Despite the most dominant post-fire fungus being a Basidiomycete, overall the community shifted from Basidiomycete to Ascomycete dominated post-fire, with particular increases in the Pezizomycete and Eurotiomycete lineages (Figure 6). The shift from Basidiomycete to Ascomycete dominance is well established in the post-fire fungal literature (Cairney & Bastias, 2007) with similar findings across diverse biomes ranging from Mediterranean shrublands of Spain (Pérez-Valera et al., 2018) to pine savannas of the American southeast (Semenova-Nelsen et al., 2019). Four of the five fire indicator taxa (Table S6) belong to the Pyronemataceae, which have long been known to fruit extensively after fires (Seaver, 1909). In addition to evidence from fruiting body surveys, the Pyronemataceae genus *Pyronema* can also dominate belowground mycelium by as much as 60% after experimental laboratory fires (Bruns et al., 2020) and increased 100 fold in frequency after prescribed fires in a *P. ponderosa* forest (Reazin et al., 2016). *Geopyxis alpina* was also an indicator of fire, and at least two species of *Geopyxis* species are known to fruit exclusively on burned soil (Wang et al., 2016). The Pyronemataceae produce sporocarps that are small orange cup fungi that appear on pyrolyzed material after burning (El-Abyad & Webster, 1968). It is possible that their color or morphology makes them better adapted to post-fire scenarios, for example many of them are orange and contain carotenoids (Carlile & Friend, 1956) which may add UV protection (Luque et al., 2012). However, it is also likely that they survive fires due to their ability to form resistant propagules called sclerotia (Smith et al., 2015) which may be thermotolerant (Richter & Barnard, 2002).

While Eurotiomycetes are widespread fungi, it is possible that their ability to grow fast and produce ubiquitous spores help them colonize open niche space after fires, as they were similarly common after wildfires in a Canadian boreal forest (Whitman et al., 2019) and in northeastern China (Yang et al., 2020). This would be analogous to the fast-colonization trait of fire adapted plants sprouting in the newly opened niche of burn zones (Keeley et al., 2011). Moreover, several species of *Aspergillus* and *Penicillium* have evidence of heat activated spores (Warcup & Baker, 1963). Much like our reasoning for why some bacterial clades responded better to fire, the Pezizomycetes may all share a similar, conserved response to fire that is less genetically complex, more basal, or more deeply conserved than the response exhibited by the Eurotiales, which may explain why all lineages within the Pezizomycetes responded positively to fire while not all lineages of the Eurotiomycetes did (Figure 6). When discussing the phylogenetic signal obtained in our analyses, we are aware of the limitations of ITS based phylogenies compared with phylogenies constructed involving 18S or 28S data and therefore the statistical significance portrayed in our consenTRAIT analysis (Figure 6) may not be as reliable as the analysis done with the 16S phylogeny (Figure 5). However, we feel confident in continuing to draw meaningful conclusions from our phylogenetic analysis as the observed signal (either positively or negatively responding) is tightly clustered within clades, and the positively responding taxa from which we form additional hypotheses (eg. Pyronemataceae, Eurotiales, Geminibasidiomycetes) have strong support in the literature behaving as observed. Future phylogenetic studies on pyrophilous fungi would benefit from sequencing which includes 18S or 28S information.

Critically analyzing similarities between certain groups of pyrophilous microbes has brought us to the hypothesis that there may be certain traits that characterize a microbe’s ability to thrive post-fire, which should be able to be placed into a conceptual framework (H3). These traits may be analogous to the traits exhibited by fire-tolerant plants (e.g. thick bark, production of serotinous cones, etc.), which themselves fall into fairly well defined suites (vegetative resprouters, structural resistance to fire, and rapid colonizers post-fire) (Keeley et al., 2011). We thus propose that microbes survive fire by analogous traits to plants and build from Grime’s Competitive-Stress-Ruderal (C-S-R) model (Grime, 1977), its adaptation to mycorrhizal fungi (Chagnon et al., 2013) and to microbial decomposers (Malik et al., 2020), and a recent study from a Canadian boreal forest fire (Whitman et al., 2019). Pyrophilous microbes appear to survive fire via trade-offs among microbial ability acquire post-fire resources (analogous to Grime’s C), tolerate heat or desiccation (analogous to Grime’s S), or early colonize or grow fast (analogous to Grime’s R) (Figure 7). Furthermore, as trade-offs among traits can be used to predict decomposition rates among wood decomposer fungi (Lustenhouwer et al., 2020), we predict that trade-offs among fire response traits might enable prediction of post-fire biogeochemical cycling. For example, the DEMENT model uses traits to improve forecasting of soil carbon by considering trade-offs between a microorganism’s ability to decompose litter versus tolerate drought (Allison & Goulden, 2017). Similarly, trade-offs between a microbe’s ability to survive fire by thermotolerance versus decomposing pyrogenic organic matter might predict changes in carbon cycling post-fire. This could be further highlighted by a trade-off of post-fire nutrient acquisition strategy microbes growing slowly as they work to breakdown recalcitrant carbon forms present in PyOM, slowly releasing this carbon for cycling, as opposed to fast-growing microbes taking advantage of easily accessible labile carbon. We hypothesize that several known pyrophilous taxa observed in this study and in others would fall into these suites of traits. For post-fire resource acquisition, basidiomycetes such as *Pholiota highlandensis* or *Lyophyllum atratum* might be more likely to degrade pyrolyzed organic matter due to their enzymatic capabilities, as supported by recent sequencing of their genomes (Steindorff et al., 2020), and *Fusarium* may be capable of denitrifying highly abundant nitrogen (Maeda et al., 2015). For thermotolerance, *Neurospora crassa* has heat-activated spores and *Rhizopogon olivaceotinctus* has spores that increase after heat treatment (Bruns et al., 2019; Emerson, 1948) and *Pyronema* may produce thermotolerant sclerotia (Moore, 1962). *Aspergillus* and *Penicillium* are likely fast-colonizers due to highly abundant spore production. However, it is also possible that certain species fall along a continuum of traits and that trade-offs may not be so strict. For example, but *Aspergillus* and *Penicillium* species are also likely thermotolerant (Warcup & Baker, 1963), and some *Penicillium* species have been found to degrade polycyclic aromatic hydrocarbons (PAHs), such as those that might be produced through combustion (Leitão, 2009). Similarly, the yeast *Basidioascus* tolerates stress with its xero- and thermo-tolerance but as a yeast may also be a fast-colonizer since it is unicellular. As our hypothesis regarding post-fire microbial trait suites expands, it may evolve into a more multi-faceted niche space as is currently used in plant trait models (Díaz et al., 2016).

**Figure 7:**
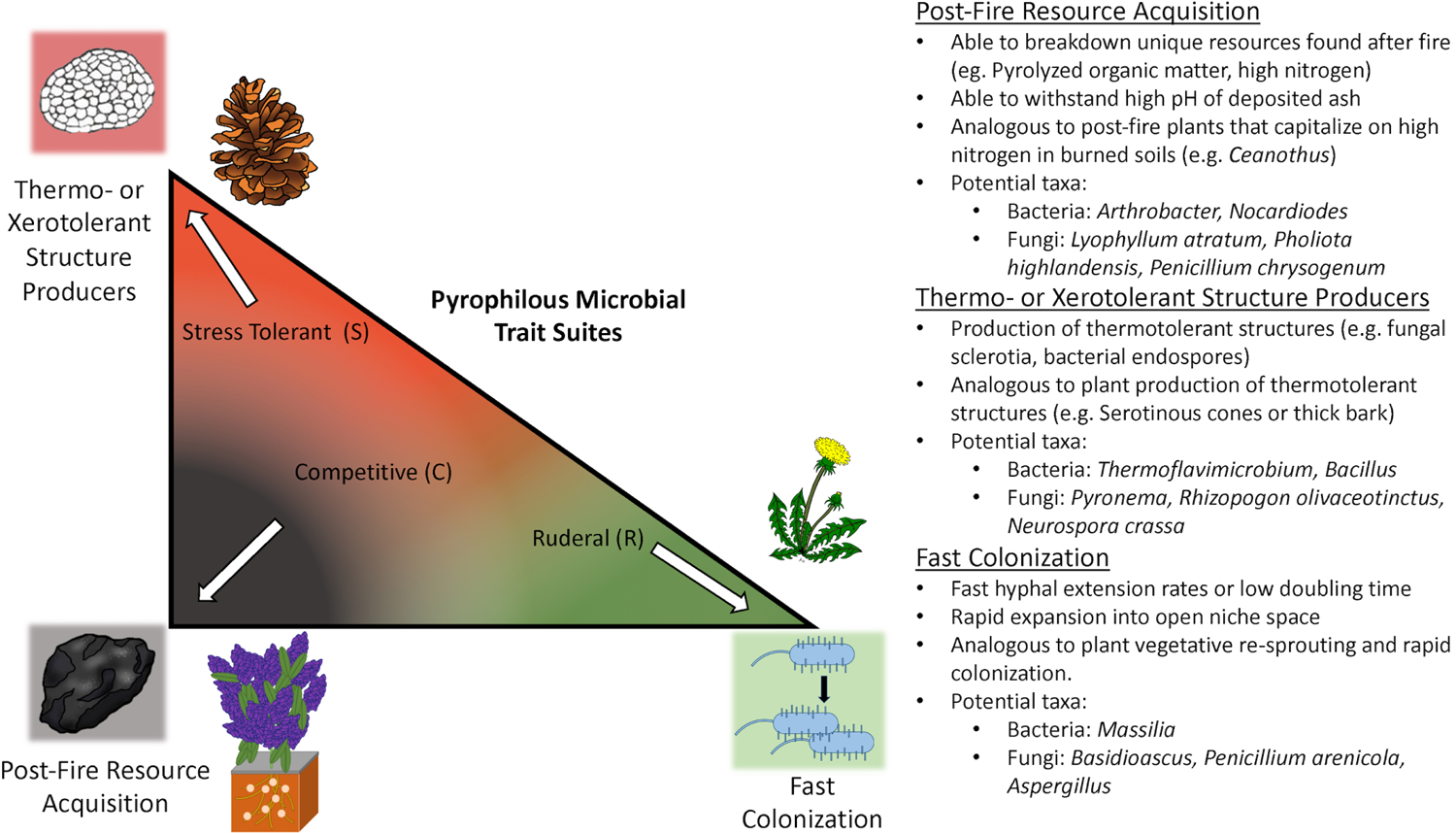
Conceptual model of hypothesized trait suites for post fire microbes, in comparison to analogous trait suits of fire adaptation in plants. Grimes CSR ecological trade-off directions are placed inside the triangle following where the corresponding pyrophilous microbial trait suite is hypothesized to relate. A color gradient between trait suites represents that these suites may not be exclusive but rather taxa may fall along a gradient between trait suites. A representation of the microbial trait is placed at each corner (Fast colonization = replicating bacteria, Thermotolerance = fungal sclerotium, Post-fire nutrient acquisition = colonized charcoal) along with an analogous plant trait representation (fast herbaceous growth, serotinous cone, root nodule forming post-fire colonizer, respectively). A brief description of the pyrophilous trait suites and hypothesized taxa placement is located on the right side of the figure.

We propose that this trait based conceptual model works for pyrophilous bacteria as well as fungi (Figure 7). For example, for post-fire resource acquisition, there are a few studies claiming *Arthrobacter* may be capable of degrading pyrogenic organic matter (Fernández-González et al., 2017; Woolet & Whitman, 2020) and so may *Nocardiodes* (Woolet & Whitman, 2020). It is also possible that ammonifying bacteria may be able to capitalize on post-fire high nitrogen content (Johnson & Curtis, 2001), and indeed one study did find that ammonifying bacteria increased post-fire while ammonia-oxidizing bacteria did not (Acea & Carballas, 1996). For thermotolerance, members of the Firmicutes are good candidates since they are almost all spore-forming bacteria (Barberán et al., 2017; Filippidou et al., 2016). For fast growth or colonization, the Proteobacteria genus *Massilia* has been hypothesized as a fast responder due to a predicted high 16S copy number (Nemergut et al., 2016; Whitman et al., 2019). We believe that categorizing pyrophilous microbes into traits will improve prediction of biogeochemical cycling post-fire. Just as redwoods survive fire with thick bark and tanoaks are vegetative resprouters, it is likely that microbes have similar strategies and this diversity of fire-responsive traits both above and belowground will enhance forest resilience to the unprecedented rise of mega-fires. However, whether they will recover from synergistic effects of introduced pathogens and global change (Simler et al., 2018) remains unclear.

## Conclusions

In conclusion, we present the first study examining the immediate effects of a megafire on both bacterial and fungal communities with a rare data set of pre- and post-fire samples and also the first study to comprehensively characterize the soil microbial communities of a redwood tanoak forest. We know of only two other instances where pre- and post-fire samples from the same locations exist (Baar et al., 1999; Glassman et al., 2016), but both were in pine forests, focused only on EMF communities, and both lacked an unburned control. We identified a massive increase in the Basidiomycete yeast *Basidioascus* and the bacterial Firmicutes post-fire, and we showed that pyrophilous bacteria and fungi and their traits are phylogenetically conserved at the class level. By comparing our work to other recent molecular characterizations of post-fire microbes in Pinaceae forests (Bruns et al., 2020; Glassman et al., 2016; Whitman et al., 2019), we can now begin to generalize traits of post-fire microbes to other forest types and compare them to analogous traits in plants (Figure 7). For example, we hypothesize certain bacteria (Firmicutes) and fungi (Pyronemataceae) appear to be able to survive fire with thermotolerant structures, and other fungi (*Penicillium*) or bacteria (*Massillia)* are fast-responders, and trade-offs might exist among these traits. Future studies of post-fire systems in a broad variety of ecosystems, and experimental determinations of microbial traits, will allow us to further characterize and generalize traits of post-fire microbes so that we can refine our conceptual model and reach the level of knowledge of post-fire traits of plants.

## Acknowledgements

We would like to acknowledge Monterey Peninsula Regional Park District (Plots 601, 603) and the University of California’s Landels-Hill Big Creek Reserve (Plot 058). We thank Michael Ernandes and Tom Bruns for their assistance in collecting pre-fire soils, Judy Chung for assistance with molecular work, and James Randolph for assistance with conceptual figure artwork. DJE is supported by the National Science Foundation Graduate Research Fellowship and the Oregon Mycological Society Scholarship.

