## Supplemental Material for "Mega-fire in Redwood Tanoak Forest Reduces Bacterial and Fungal Richness and Selects for Pyrophilous Taxa and Traits that are Phylogenetically Conserved"

**Supplementary Materials**

**Table S1**
Plot information for each of the three plots sampled before and after the Soberanes fire. All plots are in UTM Zone 10.

| Plot | Slope | Aspect | Elevation | Distance to Coast | Land Ownership |
|---|---|---|---|---|---|
| 58 | 19° | 180 | 744m | 4.1km | University of California Landels-Hill Big Creek Reserve |
| 601 | 34° | 62 | 603m | 6.4km | Monterey Peninsula Regional Park District |
| 603 | 31° | 116 | 491m | 6.3km | Monterey Peninsula Regional Park District |

**Table S2.** Summary of mean per-sample richness estimated by observed species, Shannon and Simpson diversity indices, and percent change in each plot before and after the Soberanes Fire.

**Bacteria**

| Plots | Burn status | Pre-fire | Post-fire | Percent change | P value |
|---|---|---|---|---|---|
| | Mean Per-Sample richness | | | | |
| Plots | Burn status | Pre-fire | Post-fire | Percent change | P value |
| 58 | unburned | 2,183 | 2,188 | 2.0 | 0.30 |
| 601 | burned | 2,082 | 1,255 | 40.1 | <0.01 |
| 603 | burned | 2,188 | 843 | 52.2 | <0.01 |
| | Mean Per-Sample Shannon Diversity | | | | |
| Plots | Burn status | Pre-fire | Post-fire | Percent change | P value |
| 58 | unburned | 775 | 742 | 4.3 | 0.12 |
| 601 | burned | 763 | 264 | 65.4 | <0.001 |
| 603 | burned | 782 | 171 | 78.2 | <0.001 |
| | Mean Per-Sample Simpson Diversity$^{-1}$ | | | | |
| Plots | Burn status | Pre-fire | Post-fire | Percent change | P value |
| 58 | unburned | 0.0037 | 0.005 | 18.0 | 0.49 |
| 601 | burned | 0.0034 | 0.014 | 75.7 | <0.001 |
| 603 | burned | 0.0038 | 0.021 | 81.6 | <0.001 |

**Fungi**

| Plots | Burn status | Pre-fire | Post-fire | Percent change | P value |
|---|---|---|---|---|---|
| | Mean Per-Sample Richness | | | | |
| Plots | Burn status | Pre-fire | Post-fire | Percent change | P value |
| 58 | unburned | 267 | 286 | 0.1 | 0.78 |
| 601 | burned | 229 | 96 | 37.6 | <0.01 |
| 603 | burned | 321 | 143 | 70.0 | <0.01 |
| | Mean Per-Sample Shannon Diversity | | | | |
| Plots | Burn status | Pre-fire | Post-fire | Percent change | P value |
| 58 | unburned | 39 | 42.01 | 4.7 | 0.87 |
| 601 | burned | 42 | 7.31 | 82.7 | <0.001 |
| 603 | burned | 46 | 8.46 | 81.6 | <0.001 |
| | Mean Per-Sample Simpson Diversity$^{-1}$ | | | | |
| Plots | Burn status | Pre-fire | Post-fire | Percent change | P value |
| 58 | unburned | 0.059 | 0.05 | 10.5 | 0.70 |
| 601 | burned | 0.056 | 0.26 | 78.3 | <0.001 |
| 603 | burned | 0.059 | 0.20 | 70.5 | <0.001 |

**Table S3**

Table showing all bacterial OTUs (with taxonomic identification) with at least 1% sequence abundance either pre or post-fire in burned plots (Plots 601 and 603). Percent sequences refers to the percentage of total pre-fire or post-fire sequences that OTU utilized.

| ID | Phylum | Genus | Species | % Sequences | Fire |
|---|---|---|---|---|---|
| Otu5 | Proteobacteria | *Bradyrhizobium* | *spp.* | 2.3 | Pre-fire |
| Otu14 | Proteobacteria | *Rhodoplanes* | *spp.* | 1.4 | Pre-fire |
| Otu18 | Proteobacteria | *Betaproteobacteria spp.* | *spp.* | 1.3 | Pre-fire |
| Otu24 | Proteobacteria | *Sinobacteraceae* | *spp.* | 1.2 | Pre-fire |
| Otu15 | Acidobacteria | *Acidobacteria spp.* | *spp.* | 1.1 | Pre-fire |
| Otu6 | Firmicutes | *Sporosarcina* | *spp.* | 6.9 | Post-fire |
| Otu7 | Firmicutes | *Sporosarcina* | *spp.* | 6.9 | Post-fire |
| Otu8 | Firmicutes | *Sporosarcina* | *spp.* | 6.9 | Post-fire |
| Otu9 | Actinobacteria | *Micromonospora* | *spp.* | 2.7 | Post-fire |
| Otu13 | Firmicutes | *Bacillus* | *spp.* | 1.9 | Post-fire |
| Otu1421 | Actinobacteria | *Micromonosporaceae* | *spp.* | 1.2 | Post-fire |
| Otu17024 | Firmicutes | *Bacillus* | *muralis* | 1.1 | Post-fire |
| Otu2032 | Firmicutes | *Solibacillus* | *spp.* | 1.1 | Post-fire |
| Otu61 | Actinobacteria | *Pseudonocardia* | *spp.* | 1.1 | Post-fire |
| Otu49 | Firmicutes | *Cohnella* | *spp.* | 1.1 | Post-fire |

**Table S4**

Table summarizing which fungal OTUs (with taxonomic identification) utilized at least 1% of the pre-fire or post-fire sequence abundance in the burned plots (Plots 601 and 603). Percent sequences refers to the percentage of total pre-fire or post-fire sequences that OTU utilized.

| ID | Phylum | Genus | Species | % Sequences | Fire |
|---|---|---|---|---|---|
| Otu13 | *Basidiomycota* | *Hygrocybe* | *acutoconica var. microspora* | 13.0 | Pre-fire |
| Otu23 | *Mucoromycota* | *Mortierella* | *bainieri* | 10.7 | Pre-fire |
| Otu57 | *Basidiomycota* | *Inocybe* | *flocculosa* | 3.5 | Pre-fire |
| Otu40 | *Basidiomycota* | *Cystolepiota* | *bucknallii* | 1.8 | Pre-fire |
| Otu32 | *Ascomycota* | *Cladophialophora* | *spp.* | 1.8 | Pre-fire |
| Otu89 | *Basidiomycota* | *Xeromphalina* | *cauticinalis* | 1.8 | Pre-fire |
| Otu78 | *Basidiomycota* | *Protomerulius* | *spp.* | 1.6 | Pre-fire |
| Otu63 | *Basidiomycota* | *Tomentella* | *fuscocinerea* | 1.3 | Pre-fire |
| Otu41 | *Unidentified* | *Fungus Spp.* | *spp.* | 1.2 | Pre-fire |
| Otu53 | *Ascomycota* | *Polyphilus* | *spp.* | 1.1 | Pre-fire |
| Otu49 | *Ascomycota* | *Cylindrodendrum* | *spp.* | 1.1 | Pre-fire |
| Otu156 | *Basidiomycota* | *Lyophyllum* | *semitale* | 1.0 | Pre-fire |
| Otu70 | *Basidiomycota* | *Clavaria* | *fragilis* | 1.0 | Pre-fire |
| Otu100 | *Basidiomycota* | *Ramariopsis* | *spp.* | 1.0 | Pre-fire |
| Otu1 | *Basidiomycota* | *Basidioascus* | *undulatus* | 28.2 | Post-fire |
| Otu2 | *Ascomycota* | *Tricharina* | *spp.* | 13.6 | Post-fire |
| Otu6 | *Ascomycota* | *Penicillium* | *decumbens* | 13.1 | Post-fire |
| Otu8 | *Ascomycota* | *Peziza* | *spp.* | 10.2 | Post-fire |
| Otu4 | *Ascomycota* | *Anthracobia* | *spp.* | 8.8 | Post-fire |
| Otu5 | *Ascomycota* | *Scutellinia* | *vitreola* | 7.2 | Post-fire |
| Otu7 | *Ascomycota* | *Pyronema* | *domesticum* | 4.2 | Post-fire |
| Otu16 | *Basidiomycota* | *Lyophyllum* | *semitale* | 2.6 | Post-fire |
| Otu20 | *Ascomycota* | *Fusarium* | *acutatum* | 1.2 | Post-fire |

**Table S5**
Bacterial Indicator Species Analysis results. For pre-fire bacteria the top 10 taxa are listed and identified (out of 86 identified indicators).

**Pre-fire**

| ID | p Value | Phylum | Best Taxonomic ID |
|---|---|---|---|
| Otu147 | 0.001 | Proteobacteria | Nitrosomonadales |
| Otu99 | 0.001 | Bacteroidetes | Cytophagaceae |
| Otu215 | 0.001 | Actinobacteria | Gaiellaceae |
| Otu107 | 0.001 | Proteobacteria | Sinobacteraceae |
| Otu51 | 0.001 | Proteobacteria | Rhodospirillaceae |
| Otu13706 | 0.001 | Proteobacteria | Kaistobacter |
| Otu273 | 0.001 | Acidobacteria | Acidobacteria |
| Otu13215 | 0.001 | Chloroflexi | Chloroflexi |
| Otu962 | 0.001 | Verrucomicrobia | Verrucomicrobia |
| Otu355 | 0.001 | Proteobacteria | Betaproteobacteria |

**Post-fire**

| ID | p Value | Phylum | Best Taxonomic ID |
|---|---|---|---|
| Otu84 | 0.001 | Firmicutes | Thermoactinomyceteae |
| Otu4 | 0.001 | Firmicutes | *Thermoflavimicrobium* |
| Otu2 | 0.001 | Firmicutes | *Fictibacillus* |
| Otu34 | 0.001 | Actinobacteria | Thermomonosporaceae |
| Otu12829 | 0.001 | Firmicutes | *Cohnella* |
| Otu40 | 0.001 | Firmicutes | *Tepidibacterium infernus* |
| Otu3319 | 0.001 | Firmicutes | *Paenibacillus* |
| Otu1987 | 0.001 | Actinobacteria | *Streptomyces* (Actinobacteria) |
| Otu3360 | 0.001 | Firmicutes | *Bacillus* |
| Otu33 | 0.001 | Firmicutes | Thermoactinomyceteae |
| Otu840 | 0.001 | Firmicutes | *Lysinibacillus massiliensis* |
| Otu2099 | 0.001 | Actinobacteria | Micromonosporaceae |
| Otu209 | 0.001 | Firmicutes | *Paenibacillus sp.* |
| Otu1104 | 0.001 | Firmicutes | *Paenibacillus sp.* |
| Otu689 | 0.001 | Actinobacteria | Gaiellaceae |
| Otu87 | 0.001 | Firmicutes | *Paenibacillus* |
| Otu122 | 0.001 | Proteobacteria | Betaproteobacteria |
| Otu6430 | 0.001 | Firmicutes | *Paenibacillus* |
| Otu215 | 0.001 | Actinobacteria | Gaiellaceae |
| Otu13215 | 0.001 | Chloroflexi | Chloroflexi |
| Otu1713 | 0.001 | Firmicutes | *Aneurinibacillus* |

**Table S6**
Pre-fire fungal Indicator Species Analysis results.

**Pre-fire**

| ID | p Value | Phylum | Best Taxonomic ID |
|---|---|---|---|
| Otu60 | 0.001 | Mucuromycota | *Mortierella horticola* |
| Otu23 | 0.001 | Mucuromycota | *Mortierella baineri* |
| Otu53 | 0.001 | Ascomycota | *Polyphilus frankenii* |
| Otu433 | 0.001 | Ascomycota | *Cladophialophora sp.* |
| Otu113 | 0.001 | Ascomycota | *Pycnopeziza sympodialis* |
| Otu3750 | 0.001 | Mucuromycota | *Mortierella amoeboidea* |
| Otu424 | 0.001 | Ascomycota | *Cladiophialophora sp.* |
| Otu432 | 0.001 | Ascomycota | *Thelonectria nodosa* |
| Otu597 | 0.001 | Ascomycota | *Pectenia plumbea* |
| Otu3759 | 0.001 | Mucuromycota | *Mortierella elongata* |
| Otu772 | 0.001 | Ascomycota | Pezizamycotina (sub-phylum) |
| Otu487 | 0.001 | Ascomycota | *Phomopsis sp.* |
| Otu275 | 0.001 | Ascomycota | Helotiales |

**Post-fire**

| ID | p Value | Phylum | Best Taxonomic ID |
|---|---|---|---|
| Otu2 | 0.001 | Ascomycota | *Tricharina praecox* |
| Otu8 | 0.001 | Ascomycota | *Peziza vacinii* |
| Otu4 | 0.001 | Ascomycota | *Anthracobia sp.* |
| Otu6 | 0.001 | Ascomycota | *Penicillium decumbens* |
| Otu180 | 0.001 | Ascomycota | *Geopyxis alpina* |

**Table S7**
All significant bacterial interactions to fire as reported by DeSEQ2 analysis. Bacterial taxonomic groupings at phylum or order taxonomic levels whose response was more positive (blue) or negative (orange) than expected by chance (two-tailed exact test; $P < 0.05$). "Increase" and "Decrease" refer to the number of OTUs that changed significantly and "p Value" refers to the results of the two-tailed exact test.

| Phylum | Order | Increase | Decrease | p Value | Stat |
|---|---|---|---|---|---|
| Acidobacteria | - | 6 | 0 | 0.031 | Increase |
| Acidobacteria | unidentified | 6 | 0 | 0.031 | Increase |
| Acidobacteria | unidentified | 6 | 0 | 0.031 | Increase |
| Acidobacteria | unidentified | 6 | 0 | 0.031 | Increase |
| Actinobacteria | Actinomycetales | 6 | 0 | 0.031 | Increase |
| Actinobacteria | Actinomycetales | 6 | 0 | 0.031 | Increase |
| Actinobacteria | Gaiellales | 61 | 31 | 0.002 | Increase |
| Actinobacteria | Gaiellales | 54 | 19 | 0.000 | Increase |
| Actinobacteria | Gaiellales | 54 | 19 | 0.000 | Increase |
| Firmicutes | - | 127 | 2 | 0.000 | Increase |
| Firmicutes | Bacillales | 125 | 2 | 0.000 | Increase |
| Firmicutes | Bacillales | 11 | 0 | 0.001 | Increase |
| Firmicutes | Bacillales | 12 | 1 | 0.003 | Increase |
| Firmicutes | Bacillales | 78 | 1 | 0.000 | Increase |
| Firmicutes | Bacillales | 6 | 0 | 0.031 | Increase |
| Firmicutes | Bacillales | 15 | 0 | 0.000 | Increase |
| Firmicutes | Bacillales | 10 | 0 | 0.002 | Increase |
| Firmicutes | Bacillales | 9 | 1 | 0.021 | Increase |
| Firmicutes | Bacillales | 61 | 1 | 0.000 | Increase |
| Firmicutes | Bacillales | 9 | 0 | 0.004 | Increase |
| Firmicutes | Bacillales | 8 | 0 | 0.008 | Increase |
| Firmicutes | - | 13 | 1 | 0.002 | Increase |
| Firmicutes | Clostridiales | 13 | 1 | 0.002 | Increase |
| Acidobacteria | - | 16 | 97 | 0.000 | Decrease |
| Acidobacteria | 24-Nov | 0 | 16 | 0.000 | Decrease |
| Acidobacteria | DS-100 | 0 | 10 | 0.002 | Decrease |
| Acidobacteria | RB41 | 10 | 55 | 0.000 | Decrease |
| Acidobacteria | unidentified | 1 | 9 | 0.021 | Decrease |
| Acidobacteria | 24-Nov | 0 | 16 | 0.000 | Decrease |
| Acidobacteria | DS-100 | 0 | 10 | 0.002 | Decrease |
| Acidobacteria | RB41 | 7 | 22 | 0.008 | Decrease |
| Acidobacteria | RB41 | 3 | 33 | 0.000 | Decrease |
| Acidobacteria | unidentified | 1 | 9 | 0.021 | Decrease |
| Acidobacteria | 24-Nov | 0 | 16 | 0.000 | Decrease |

| | | | | | |
|---|---|---|---|---|---|
| Acidobacteria | DS-100 | 0 | 10 | 0.002 | Decrease |
| Acidobacteria | RB41 | 7 | 22 | 0.008 | Decrease |
| Acidobacteria | RB41 | 3 | 33 | 0.000 | Decrease |
| Acidobacteria | unidentified | 1 | 9 | 0.021 | Decrease |
| Acidobacteria | - | 5 | 19 | 0.007 | Decrease |
| Acidobacteria | unidentified | 5 | 19 | 0.007 | Decrease |
| Acidobacteria | unidentified | 5 | 19 | 0.007 | Decrease |
| Acidobacteria | unidentified | 5 | 19 | 0.007 | Decrease |
| Acidobacteria | - | 55 | 89 | 0.006 | Decrease |
| Acidobacteria | iii1-15 | 43 | 79 | 0.001 | Decrease |
| Acidobacteria | iii1-15 | 27 | 62 | 0.000 | Decrease |
| Acidobacteria | iii1-15 | 27 | 62 | 0.000 | Decrease |
| Acidobacteria | - | 6 | 24 | 0.001 | Decrease |
| Acidobacteria | Acidobacteriales | 6 | 24 | 0.001 | Decrease |
| Acidobacteria | Acidobacteriales | 4 | 17 | 0.007 | Decrease |
| Acidobacteria | Acidobacteriales | 2 | 11 | 0.022 | Decrease |
| Acidobacteria | - | 3 | 18 | 0.001 | Decrease |
| Acidobacteria | DS-18 | 1 | 16 | 0.000 | Decrease |
| Acidobacteria | DS-18 | 1 | 16 | 0.000 | Decrease |
| Acidobacteria | DS-18 | 1 | 16 | 0.000 | Decrease |
| Acidobacteria | - | 10 | 72 | 0.000 | Decrease |
| Acidobacteria | Solibacterales | 10 | 72 | 0.000 | Decrease |
| Acidobacteria | Solibacterales | 2 | 25 | 0.000 | Decrease |
| Acidobacteria | Solibacterales | 7 | 46 | 0.000 | Decrease |
| Acidobacteria | Solibacterales | 1 | 20 | 0.000 | Decrease |
| Acidobacteria | Solibacterales | 7 | 46 | 0.000 | Decrease |
| Actinobacteria | Solirubrobacterales | 5 | 17 | 0.017 | Decrease |
| Actinobacteria | Solirubrobacterales | 4 | 16 | 0.012 | Decrease |
| Armatimonadetes | - | 1 | 16 | 0.000 | Decrease |
| Armatimonadetes | [Fimbriimonadales] | 1 | 16 | 0.000 | Decrease |
| Armatimonadetes | [Fimbriimonadales] | 1 | 15 | 0.001 | Decrease |
| Armatimonadetes | [Fimbriimonadales] | 1 | 13 | 0.002 | Decrease |
| Armatimonadetes | - | 5 | 26 | 0.000 | Decrease |
| Armatimonadetes | unidentified | 5 | 26 | 0.000 | Decrease |
| Armatimonadetes | unidentified | 5 | 26 | 0.000 | Decrease |
| Armatimonadetes | unidentified | 5 | 26 | 0.000 | Decrease |
| Armatimonadetes | - | 0 | 6 | 0.031 | Decrease |
| Armatimonadetes | - | 1 | 9 | 0.021 | Decrease |
| Armatimonadetes | SJA-22 | 1 | 8 | 0.039 | Decrease |
| Armatimonadetes | SJA-22 | 1 | 8 | 0.039 | Decrease |
| Armatimonadetes | SJA-22 | 1 | 8 | 0.039 | Decrease |
| Bacteroidetes | - | 26 | 179 | 0.000 | Decrease |

| | | | | | |
|---|---|---|---|---|---|
| Bacteroidetes | [Saprospirales] | 26 | 179 | 0.000 | Decrease |
| Bacteroidetes | [Saprospirales] | 20 | 164 | 0.000 | Decrease |
| Bacteroidetes | [Saprospirales] | 2 | 10 | 0.039 | Decrease |
| Bacteroidetes | [Saprospirales] | 0 | 9 | 0.004 | Decrease |
| Bacteroidetes | [Saprospirales] | 18 | 137 | 0.000 | Decrease |
| Bacteroidetes | - | 11 | 75 | 0.000 | Decrease |
| Bacteroidetes | Cytophagales | 11 | 75 | 0.000 | Decrease |
| Bacteroidetes | Cytophagales | 11 | 72 | 0.000 | Decrease |
| Bacteroidetes | Cytophagales | 9 | 58 | 0.000 | Decrease |
| Bacteroidetes | - | 5 | 19 | 0.007 | Decrease |
| Bacteroidetes | Flavobacteriales | 5 | 19 | 0.007 | Decrease |
| Bacteroidetes | Flavobacteriales | 3 | 12 | 0.035 | Decrease |
| Bacteroidetes | - | 11 | 100 | 0.000 | Decrease |
| Bacteroidetes | Sphingobacteriales | 11 | 100 | 0.000 | Decrease |
| Bacteroidetes | Sphingobacteriales | 1 | 14 | 0.001 | Decrease |
| Bacteroidetes | Sphingobacteriales | 10 | 86 | 0.000 | Decrease |
| Bacteroidetes | Sphingobacteriales | 1 | 13 | 0.002 | Decrease |
| Bacteroidetes | Sphingobacteriales | 10 | 86 | 0.000 | Decrease |
| Chlamydiae | - | 13 | 28 | 0.028 | Decrease |
| Chlamydiae | Chlamydiales | 12 | 27 | 0.024 | Decrease |
| Chlamydiae | Chlamydiales | 7 | 24 | 0.003 | Decrease |
| Chlorobi | - | 0 | 6 | 0.031 | Decrease |
| Chlorobi | unidentified | 0 | 6 | 0.031 | Decrease |
| Chlorobi | unidentified | 0 | 6 | 0.031 | Decrease |
| Chlorobi | unidentified | 0 | 6 | 0.031 | Decrease |
| Chloroflexi | - | 24 | 74 | 0.000 | Decrease |
| Chloroflexi | SBR1031 | 11 | 39 | 0.000 | Decrease |
| Chloroflexi | SBR1031 | 8 | 28 | 0.001 | Decrease |
| Chloroflexi | SBR1031 | 2 | 11 | 0.022 | Decrease |
| Chloroflexi | SBR1031 | 8 | 28 | 0.001 | Decrease |
| Chloroflexi | SBR1031 | 2 | 11 | 0.022 | Decrease |
| Chloroflexi | - | 6 | 23 | 0.002 | Decrease |
| Chloroflexi | Chloroflexales | 0 | 6 | 0.031 | Decrease |
| Chloroflexi | - | 7 | 39 | 0.000 | Decrease |
| Chloroflexi | AKYG885 | 1 | 19 | 0.000 | Decrease |
| Chloroflexi | unidentified | 0 | 12 | 0.000 | Decrease |
| Chloroflexi | AKYG885 | 0 | 7 | 0.016 | Decrease |
| Chloroflexi | AKYG885 | 1 | 12 | 0.003 | Decrease |
| Chloroflexi | unidentified | 0 | 12 | 0.000 | Decrease |
| Chloroflexi | AKYG885 | 0 | 7 | 0.016 | Decrease |
| Chloroflexi | AKYG885 | 1 | 12 | 0.003 | Decrease |
| Chloroflexi | unidentified | 0 | 12 | 0.000 | Decrease |

| | | | | | |
|---|---|---|---|---|---|
| Cyanobacteria | - | 1 | 11 | 0.006 | Decrease |
| Cyanobacteria | MLE1-12 | 1 | 8 | 0.039 | Decrease |
| Cyanobacteria | MLE1-12 | 1 | 8 | 0.039 | Decrease |
| Cyanobacteria | MLE1-12 | 1 | 8 | 0.039 | Decrease |
| Elusimicrobia | - | 7 | 36 | 0.000 | Decrease |
| Elusimicrobia | FAC88 | 1 | 15 | 0.001 | Decrease |
| Elusimicrobia | IIb | 2 | 14 | 0.004 | Decrease |
| Elusimicrobia | FAC88 | 1 | 15 | 0.001 | Decrease |
| Elusimicrobia | IIb | 2 | 14 | 0.004 | Decrease |
| Elusimicrobia | FAC88 | 1 | 15 | 0.001 | Decrease |
| Elusimicrobia | IIb | 2 | 14 | 0.004 | Decrease |
| Elusimicrobia | - | 0 | 9 | 0.004 | Decrease |
| Elusimicrobia | unidentified | 0 | 9 | 0.004 | Decrease |
| Elusimicrobia | unidentified | 0 | 9 | 0.004 | Decrease |
| Elusimicrobia | unidentified | 0 | 9 | 0.004 | Decrease |
| FBP | - | 0 | 8 | 0.008 | Decrease |
| FBP | unidentified | 0 | 8 | 0.008 | Decrease |
| FBP | unidentified | 0 | 8 | 0.008 | Decrease |
| FBP | unidentified | 0 | 8 | 0.008 | Decrease |
| Gemmatimonadetes | - | 10 | 28 | 0.005 | Decrease |
| Gemmatimonadetes | unidentified | 10 | 28 | 0.005 | Decrease |
| Gemmatimonadetes | unidentified | 10 | 28 | 0.005 | Decrease |
| Gemmatimonadetes | unidentified | 10 | 28 | 0.005 | Decrease |
| Gemmatimonadetes | - | 1 | 10 | 0.012 | Decrease |
| Gemmatimonadetes | unidentified | 1 | 10 | 0.012 | Decrease |
| Gemmatimonadetes | unidentified | 1 | 10 | 0.012 | Decrease |
| Gemmatimonadetes | unidentified | 1 | 10 | 0.012 | Decrease |
| Gemmatimonadetes | - | 33 | 91 | 0.000 | Decrease |
| Gemmatimonadetes | Ellin5290 | 7 | 26 | 0.001 | Decrease |
| Gemmatimonadetes | Gemmatimonadales | 5 | 23 | 0.001 | Decrease |
| Gemmatimonadetes | Ellin5290 | 7 | 26 | 0.001 | Decrease |
| Gemmatimonadetes | Gemmatimonadales | 0 | 6 | 0.031 | Decrease |
| Gemmatimonadetes | Ellin5290 | 7 | 26 | 0.001 | Decrease |
| Gemmatimonadetes | Gemmatimonadales | 0 | 6 | 0.031 | Decrease |
| OP3 | - | 3 | 23 | 0.000 | Decrease |
| OP3 | GIF10 | 1 | 10 | 0.012 | Decrease |
| OP3 | unidentified | 2 | 13 | 0.007 | Decrease |
| OP3 | GIF10 | 1 | 10 | 0.012 | Decrease |
| OP3 | unidentified | 2 | 13 | 0.007 | Decrease |
| OP3 | GIF10 | 1 | 10 | 0.012 | Decrease |
| OP3 | unidentified | 2 | 13 | 0.007 | Decrease |
| Planctomycetes | - | 33 | 169 | 0.000 | Decrease |

| | | | | | |
|---|---|---|---|---|---|
| Planctomycetes | WD2101 | 10 | 120 | 0.000 | Decrease |
| Planctomycetes | WD2101 | 10 | 120 | 0.000 | Decrease |
| Planctomycetes | WD2101 | 10 | 120 | 0.000 | Decrease |
| Planctomycetes | - | 98 | 463 | 0.000 | Decrease |
| Planctomycetes | Gemmatales | 71 | 387 | 0.000 | Decrease |
| Planctomycetes | Pirellulales | 21 | 49 | 0.001 | Decrease |
| Planctomycetes | Planctomycetales | 5 | 23 | 0.001 | Decrease |
| Planctomycetes | Gemmatales | 66 | 335 | 0.000 | Decrease |
| Planctomycetes | Gemmatales | 5 | 52 | 0.000 | Decrease |
| Planctomycetes | Pirellulales | 21 | 49 | 0.001 | Decrease |
| Planctomycetes | Planctomycetales | 5 | 23 | 0.001 | Decrease |
| Planctomycetes | Gemmatales | 8 | 76 | 0.000 | Decrease |
| Planctomycetes | Gemmatales | 58 | 259 | 0.000 | Decrease |
| Planctomycetes | Gemmatales | 5 | 51 | 0.000 | Decrease |
| Planctomycetes | Pirellulales | 19 | 38 | 0.016 | Decrease |
| Planctomycetes | Planctomycetales | 5 | 23 | 0.001 | Decrease |
| Planctomycetes | - | 0 | 19 | 0.000 | Decrease |
| Planctomycetes | DH61 | 0 | 8 | 0.008 | Decrease |
| Planctomycetes | p04_C01 | 0 | 11 | 0.001 | Decrease |
| Planctomycetes | DH61 | 0 | 8 | 0.008 | Decrease |
| Planctomycetes | p04_C01 | 0 | 11 | 0.001 | Decrease |
| Planctomycetes | DH61 | 0 | 8 | 0.008 | Decrease |
| Planctomycetes | p04_C01 | 0 | 11 | 0.001 | Decrease |
| Proteobacteria | - | 100 | 452 | 0.000 | Decrease |
| Proteobacteria | Caulobacterales | 2 | 24 | 0.000 | Decrease |
| Proteobacteria | Ellin329 | 3 | 20 | 0.000 | Decrease |
| Proteobacteria | Rhizobiales | 26 | 161 | 0.000 | Decrease |
| Proteobacteria | Rhodobacterales | 5 | 19 | 0.007 | Decrease |
| Proteobacteria | Rhodospirillales | 37 | 138 | 0.000 | Decrease |
| Proteobacteria | Sphingomonadales | 10 | 39 | 0.000 | Decrease |
| Proteobacteria | unidentified | 6 | 28 | 0.000 | Decrease |
| Proteobacteria | Caulobacterales | 2 | 24 | 0.000 | Decrease |
| Proteobacteria | Ellin329 | 3 | 20 | 0.000 | Decrease |
| Proteobacteria | Rhizobiales | 0 | 7 | 0.016 | Decrease |
| Proteobacteria | Rhizobiales | 14 | 64 | 0.000 | Decrease |
| Proteobacteria | Rhizobiales | 0 | 13 | 0.000 | Decrease |
| Proteobacteria | Rhizobiales | 0 | 12 | 0.000 | Decrease |
| Proteobacteria | Rhizobiales | 3 | 29 | 0.000 | Decrease |
| Proteobacteria | Rhizobiales | 0 | 7 | 0.016 | Decrease |
| Proteobacteria | Rhodobacterales | 4 | 15 | 0.019 | Decrease |
| Proteobacteria | Rhodospirillales | 4 | 27 | 0.000 | Decrease |
| Proteobacteria | Rhodospirillales | 30 | 104 | 0.000 | Decrease |

| | | | | | |
|---|---|---|---|---|---|
| Proteobacteria | Sphingomonadales | 8 | 36 | 0.000 | Decrease |
| Proteobacteria | unidentified | 6 | 28 | 0.000 | Decrease |
| Proteobacteria | Caulobacterales | 2 | 16 | 0.001 | Decrease |
| Proteobacteria | Ellin329 | 3 | 20 | 0.000 | Decrease |
| Proteobacteria | Rhizobiales | 0 | 7 | 0.016 | Decrease |
| Proteobacteria | Rhizobiales | 0 | 8 | 0.008 | Decrease |
| Proteobacteria | Rhizobiales | 7 | 39 | 0.000 | Decrease |
| Proteobacteria | Rhizobiales | 0 | 12 | 0.000 | Decrease |
| Proteobacteria | Rhizobiales | 0 | 6 | 0.031 | Decrease |
| Proteobacteria | Rhizobiales | 3 | 29 | 0.000 | Decrease |
| Proteobacteria | Rhodobacterales | 4 | 15 | 0.019 | Decrease |
| Proteobacteria | Rhodospirillales | 4 | 24 | 0.000 | Decrease |
| Proteobacteria | Rhodospirillales | 30 | 97 | 0.000 | Decrease |
| Proteobacteria | Sphingomonadales | 1 | 11 | 0.006 | Decrease |
| Proteobacteria | Sphingomonadales | 0 | 13 | 0.000 | Decrease |
| Proteobacteria | unidentified | 6 | 28 | 0.000 | Decrease |
| Proteobacteria | - | 47 | 133 | 0.000 | Decrease |
| Proteobacteria | Burkholderiales | 3 | 42 | 0.000 | Decrease |
| Proteobacteria | unidentified | 12 | 26 | 0.034 | Decrease |
| Proteobacteria | Burkholderiales | 0 | 20 | 0.000 | Decrease |
| Proteobacteria | Burkholderiales | 2 | 14 | 0.004 | Decrease |
| Proteobacteria | unidentified | 12 | 26 | 0.034 | Decrease |
| Proteobacteria | Burkholderiales | 0 | 11 | 0.001 | Decrease |
| Proteobacteria | Burkholderiales | 2 | 10 | 0.039 | Decrease |
| Proteobacteria | unidentified | 12 | 26 | 0.034 | Decrease |
| Proteobacteria | - | 114 | 491 | 0.000 | Decrease |
| Proteobacteria | Bdellovibrionales | 8 | 53 | 0.000 | Decrease |
| Proteobacteria | FAC87 | 0 | 10 | 0.002 | Decrease |
| Proteobacteria | MIZ46 | 6 | 50 | 0.000 | Decrease |
| Proteobacteria | Myxococcales | 65 | 276 | 0.000 | Decrease |
| Proteobacteria | Syntrophobacterales | 18 | 55 | 0.000 | Decrease |
| Proteobacteria | unidentified | 4 | 24 | 0.000 | Decrease |
| Proteobacteria | Bdellovibrionales | 4 | 50 | 0.000 | Decrease |
| Proteobacteria | FAC87 | 0 | 10 | 0.002 | Decrease |
| Proteobacteria | MIZ46 | 6 | 50 | 0.000 | Decrease |
| Proteobacteria | Myxococcales | 9 | 28 | 0.003 | Decrease |
| Proteobacteria | Myxococcales | 2 | 15 | 0.002 | Decrease |
| Proteobacteria | Myxococcales | 5 | 24 | 0.001 | Decrease |
| Proteobacteria | Myxococcales | 39 | 190 | 0.000 | Decrease |
| Proteobacteria | Syntrophobacterales | 18 | 55 | 0.000 | Decrease |
| Proteobacteria | unidentified | 4 | 24 | 0.000 | Decrease |
| Proteobacteria | Bdellovibrionales | 4 | 47 | 0.000 | Decrease |

| | | | | | |
|---|---|---|---|---|---|
| Proteobacteria | FAC87 | 0 | 10 | 0.002 | Decrease |
| Proteobacteria | MIZ46 | 6 | 50 | 0.000 | Decrease |
| Proteobacteria | Myxococcales | 8 | 28 | 0.001 | Decrease |
| Proteobacteria | Myxococcales | 1 | 11 | 0.006 | Decrease |
| Proteobacteria | Myxococcales | 3 | 21 | 0.000 | Decrease |
| Proteobacteria | Myxococcales | 39 | 190 | 0.000 | Decrease |
| Proteobacteria | Syntrophobacterales | 18 | 55 | 0.000 | Decrease |
| Proteobacteria | unidentified | 4 | 24 | 0.000 | Decrease |
| Proteobacteria | - | 39 | 124 | 0.000 | Decrease |
| Proteobacteria | Legionellales | 14 | 40 | 0.001 | Decrease |
| Proteobacteria | Xanthomonadales | 14 | 64 | 0.000 | Decrease |
| Proteobacteria | Legionellales | 6 | 26 | 0.001 | Decrease |
| Proteobacteria | Xanthomonadales | 8 | 51 | 0.000 | Decrease |
| Proteobacteria | Legionellales | 0 | 8 | 0.008 | Decrease |
| Proteobacteria | Legionellales | 6 | 18 | 0.023 | Decrease |
| Proteobacteria | Xanthomonadales | 1 | 9 | 0.021 | Decrease |
| Proteobacteria | Xanthomonadales | 7 | 41 | 0.000 | Decrease |
| TM7 | - | 0 | 10 | 0.002 | Decrease |
| TM7 | unidentified | 0 | 10 | 0.002 | Decrease |
| TM7 | unidentified | 0 | 10 | 0.002 | Decrease |
| TM7 | unidentified | 0 | 10 | 0.002 | Decrease |
| Verrucomicrobia | - | 1 | 12 | 0.003 | Decrease |
| Verrucomicrobia | S-BQ2-57 | 1 | 10 | 0.012 | Decrease |
| Verrucomicrobia | S-BQ2-57 | 1 | 10 | 0.012 | Decrease |
| Verrucomicrobia | S-BQ2-57 | 1 | 10 | 0.012 | Decrease |
| Verrucomicrobia | - | 39 | 118 | 0.000 | Decrease |
| Verrucomicrobia | [Pedosphaerales] | 39 | 117 | 0.000 | Decrease |
| Verrucomicrobia | [Pedosphaerales] | 3 | 35 | 0.000 | Decrease |
| Verrucomicrobia | [Pedosphaerales] | 20 | 48 | 0.001 | Decrease |
| Verrucomicrobia | [Pedosphaerales] | 3 | 35 | 0.000 | Decrease |
| Verrucomicrobia | [Pedosphaerales] | 20 | 48 | 0.001 | Decrease |
| Verrucomicrobia | - | 9 | 80 | 0.000 | Decrease |
| Verrucomicrobia | [Chthoniobacterales] | 9 | 80 | 0.000 | Decrease |
| Verrucomicrobia | [Chthoniobacterales] | 9 | 80 | 0.000 | Decrease |
| Verrucomicrobia | [Chthoniobacterales] | 0 | 6 | 0.031 | Decrease |
| Verrucomicrobia | [Chthoniobacterales] | 2 | 21 | 0.000 | Decrease |
| Verrucomicrobia | [Chthoniobacterales] | 0 | 14 | 0.000 | Decrease |
| Verrucomicrobia | [Chthoniobacterales] | 3 | 28 | 0.000 | Decrease |
| Verrucomicrobia | - | 1 | 18 | 0.000 | Decrease |
| Verrucomicrobia | Opitutales | 0 | 16 | 0.000 | Decrease |
| Verrucomicrobia | Opitutales | 0 | 16 | 0.000 | Decrease |
| Verrucomicrobia | Opitutales | 0 | 11 | 0.001 | Decrease |

| | | | | | |
|---|---|---|---|---|---|
| Verrucomicrobia | - | 8 | 22 | 0.016 | Decrease |
| Verrucomicrobia | Verrucomicrobiales | 8 | 22 | 0.016 | Decrease |
| Verrucomicrobia | Verrucomicrobiales | 8 | 22 | 0.016 | Decrease |
| Verrucomicrobia | Verrucomicrobiales | 6 | 19 | 0.015 | Decrease |

**Table S8**

All significant fungal interactions to fire as reported by DeSEQ2 analysis. Fungal taxonomic groupings at phylum or order taxonomic levels whose response was more positive (blue) or negative (orange) than expected by chance (two-tailed exact test; $P < 0.05$). "Increase" and "Decrease" refer to the number of OTUs that changed significantly and "p Value" refers to the results of the two-tailed exact test. (see corresponding .xlsx file)

| Phylum | Order | Increase | Decrease | p Value | Stat |
|---|---|---|---|---|---|
| Ascomycota | Eurotiales | 53 | 21 | 0.000 | Increase |
| Ascomycota | Onygenales | 14 | 3 | 0.013 | Increase |
| Ascomycota | Eurotiales | 44 | 17 | 0.001 | Increase |
| Ascomycota | Eurotiales | 31 | 13 | 0.010 | Increase |
| Ascomycota | - | 55 | 34 | 0.033 | Increase |
| Ascomycota | Pezizales | 55 | 34 | 0.033 | Increase |
| Ascomycota | Pezizales | 39 | 17 | 0.005 | Increase |
| Ascomycota | Pezizales | 16 | 3 | 0.004 | Increase |
| Basidiomycota | Russulales | 9 | 1 | 0.021 | Increase |
| Basidiomycota | Russulales | 8 | 0 | 0.008 | Increase |
| Basidiomycota | - | 24 | 6 | 0.001 | Increase |
| Basidiomycota | Geminibasidiales | 24 | 6 | 0.001 | Increase |
| Basidiomycota | Geminibasidiales | 24 | 6 | 0.001 | Increase |
| Basidiomycota | Geminibasidiales | 19 | 0 | 0.000 | Increase |
| Ascomycota | - | 103 | 158 | 0.001 | Decrease |
| Ascomycota | Capnodiales | 21 | 38 | 0.036 | Decrease |
| Ascomycota | Capnodiales | 5 | 16 | 0.027 | Decrease |
| Ascomycota | Pleosporales | 0 | 6 | 0.031 | Decrease |
| Ascomycota | Chaetothyriales | 51 | 96 | 0.000 | Decrease |
| Ascomycota | Chaetothyriales | 34 | 76 | 0.000 | Decrease |
| Ascomycota | Chaetothyriales | 9 | 36 | 0.000 | Decrease |
| Ascomycota | - | 101 | 151 | 0.002 | Decrease |
| Ascomycota | Helotiales | 95 | 130 | 0.023 | Decrease |
| Ascomycota | unidentified | 1 | 10 | 0.012 | Decrease |
| Ascomycota | Helotiales | 31 | 52 | 0.028 | Decrease |
| Ascomycota | unidentified | 1 | 10 | 0.012 | Decrease |
| Ascomycota | Helotiales | 31 | 52 | 0.028 | Decrease |
| Ascomycota | unidentified | 1 | 10 | 0.012 | Decrease |
| Ascomycota | Orbiliales | 1 | 9 | 0.021 | Decrease |
| Ascomycota | - | 20 | 37 | 0.033 | Decrease |
| Ascomycota | unidentified | 20 | 37 | 0.033 | Decrease |
| Ascomycota | unidentified | 20 | 37 | 0.033 | Decrease |
| Ascomycota | unidentified | 20 | 37 | 0.033 | Decrease |
| Basidiomycota | Agaricales | 2 | 10 | 0.039 | Decrease |

| | | | | | |
|---|---|---|---|---|---|
| Basidiomycota | Boletales | 1 | 8 | 0.039 | Decrease |
| Basidiomycota | Boletales | 1 | 8 | 0.039 | Decrease |
| Glomeromycota | - | 1 | 13 | 0.002 | Decrease |
| Glomeromycota | Glomerales | 1 | 13 | 0.002 | Decrease |
| Glomeromycota | Glomerales | 1 | 13 | 0.002 | Decrease |
| Glomeromycota | Glomerales | 0 | 9 | 0.004 | Decrease |
| Mortierellomycota | - | 37 | 60 | 0.025 | Decrease |
| Mortierellomycota | Mortierellales | 37 | 60 | 0.025 | Decrease |
| Mortierellomycota | Mortierellales | 37 | 60 | 0.025 | Decrease |
| Mortierellomycota | Mortierellales | 37 | 60 | 0.025 | Decrease |
| Mucoromycota | - | 1 | 9 | 0.021 | Decrease |
| Mucoromycota | Umbelopsidales | 1 | 9 | 0.021 | Decrease |
| Mucoromycota | Umbelopsidales | 1 | 9 | 0.021 | Decrease |
| Mucoromycota | Umbelopsidales | 1 | 9 | 0.021 | Decrease |

**Figure S1**
Plot map of plot 058, the unburned control plot.

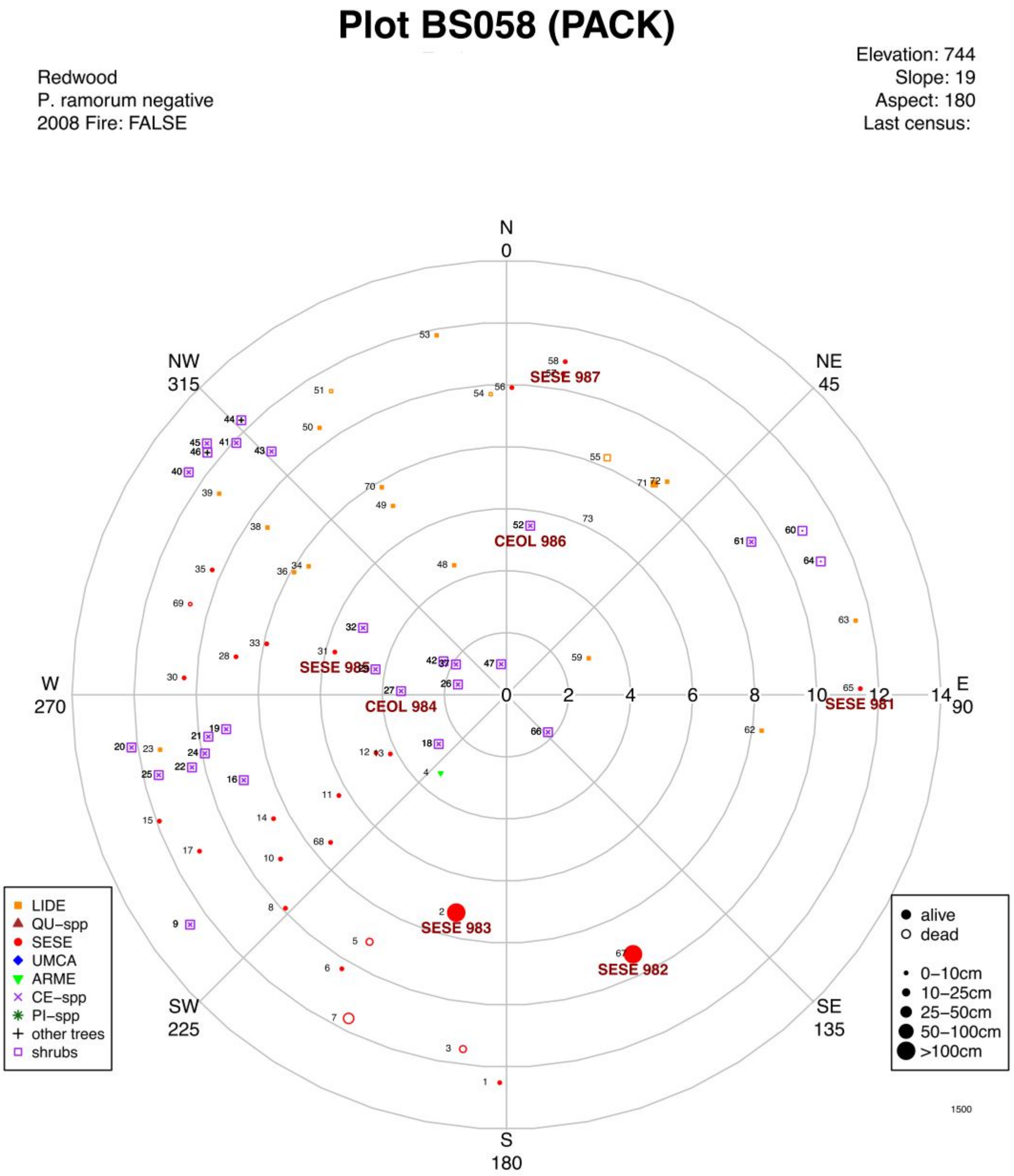

**Figure S2**
Plot map of plot 601, one of the two burned plots.

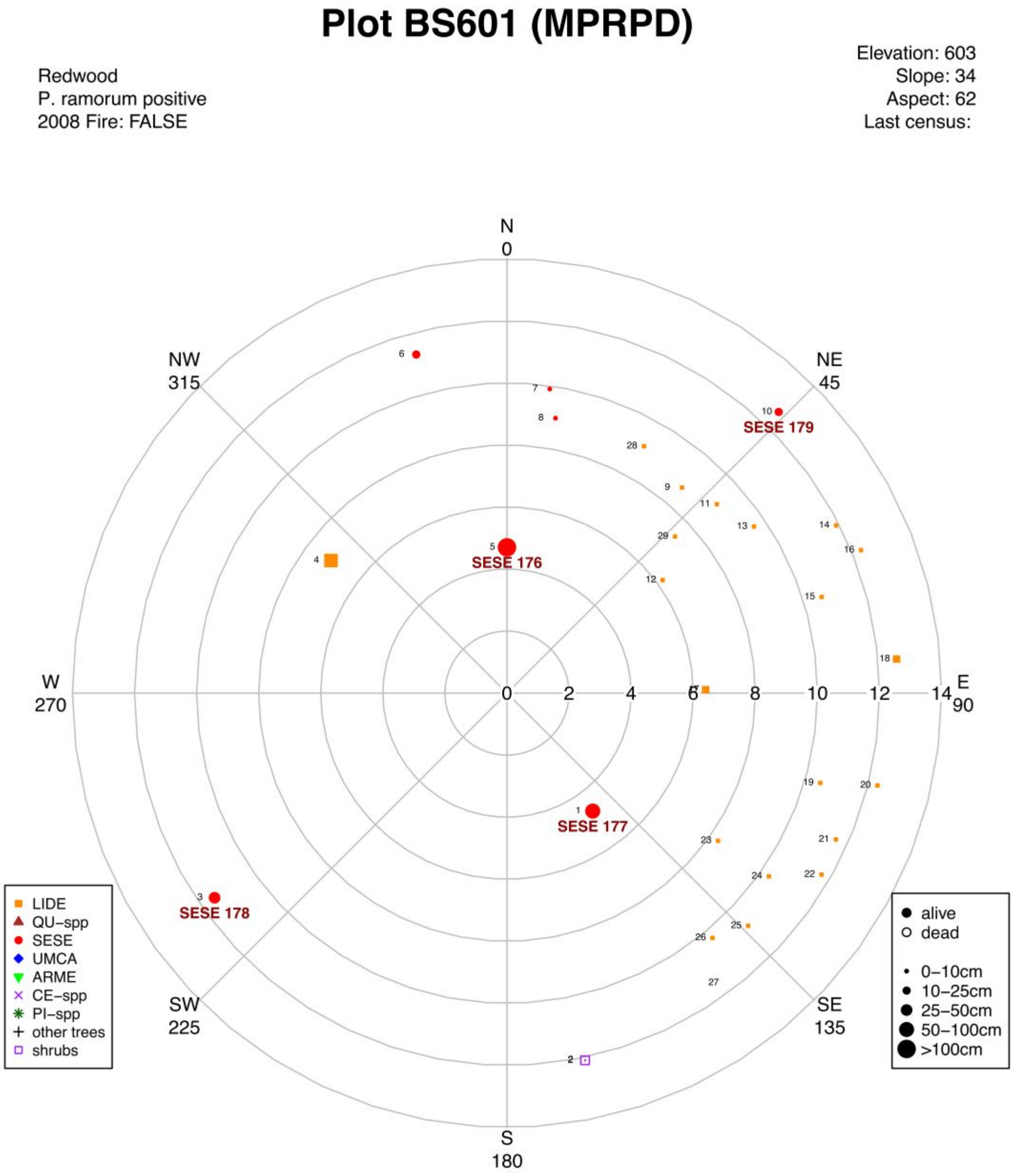

**Figure S3**
Plot map of plot 603, one of the two burned plots.

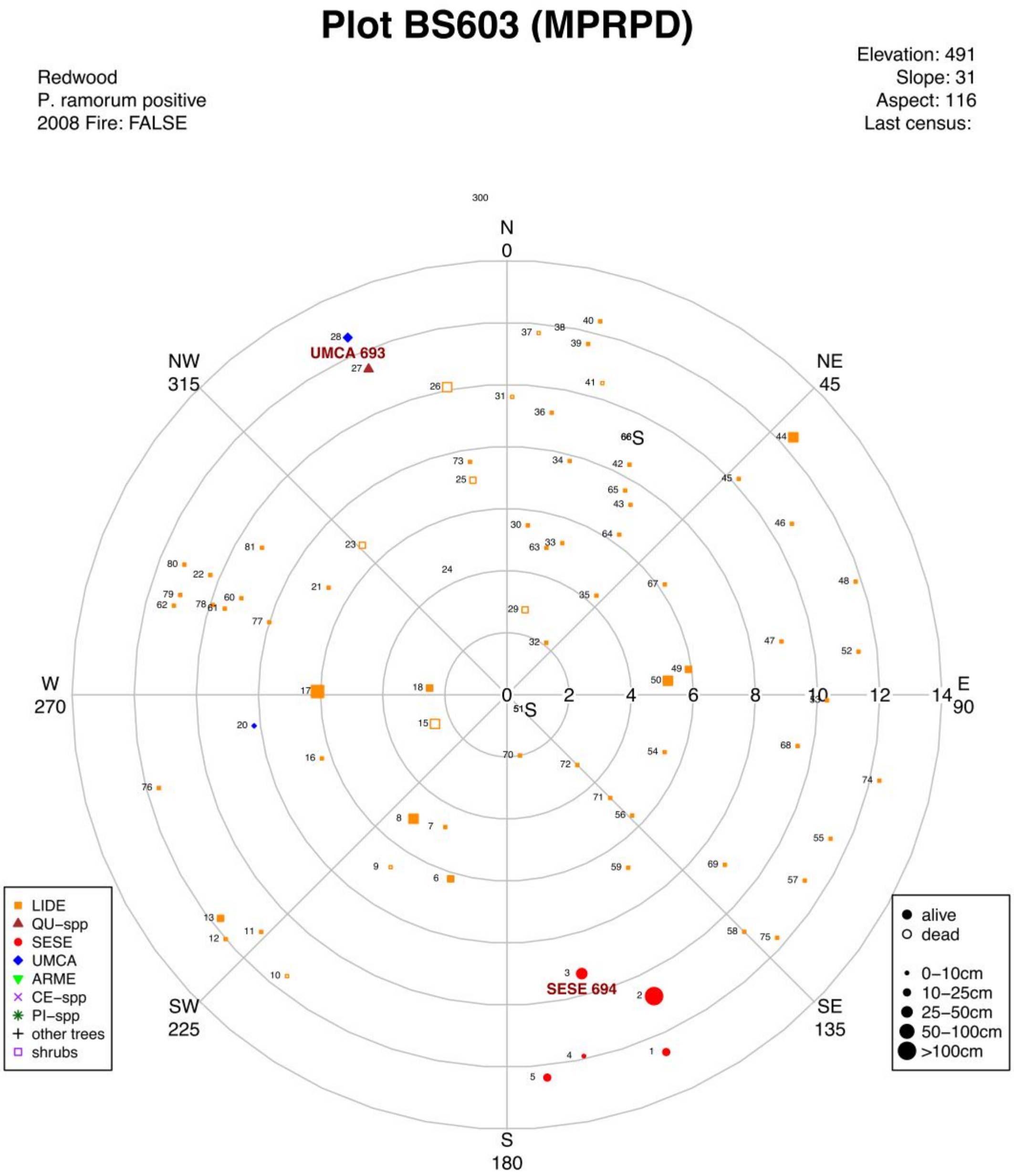

**Figure S4**

Correlation of bacterial (A & B) and fungal (C & D) species observed with ACE and Chao1 richness indices. Due to high correlation of metrics, observed species was chosen.

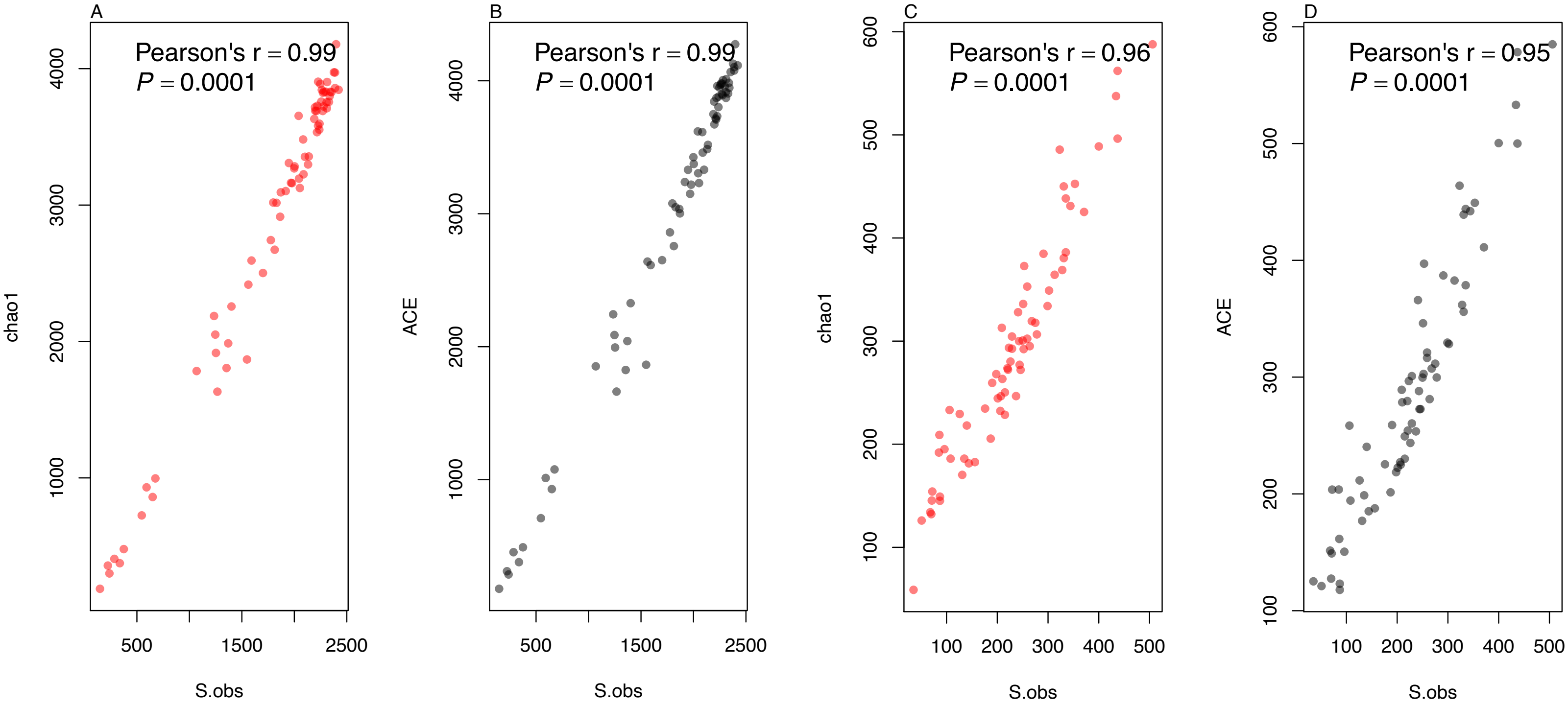

**Figure S5**
Mean per sample number of observed species for Ectomycorrhizal, Saprotrophic, and Arbuscular mycorrhizal fungi. Colors differentiate pre- and post-fire and shapes differentiate burned and unburned plots. Statistically significant difference in richness was tested using ANOVA (for burned plots, $F_{1,1} = 84.91$, $p < 0.001$, for unburned $F_{1,1} = 4.119$. $p = 0.0547$). Letters represent Tukey HSD differences.

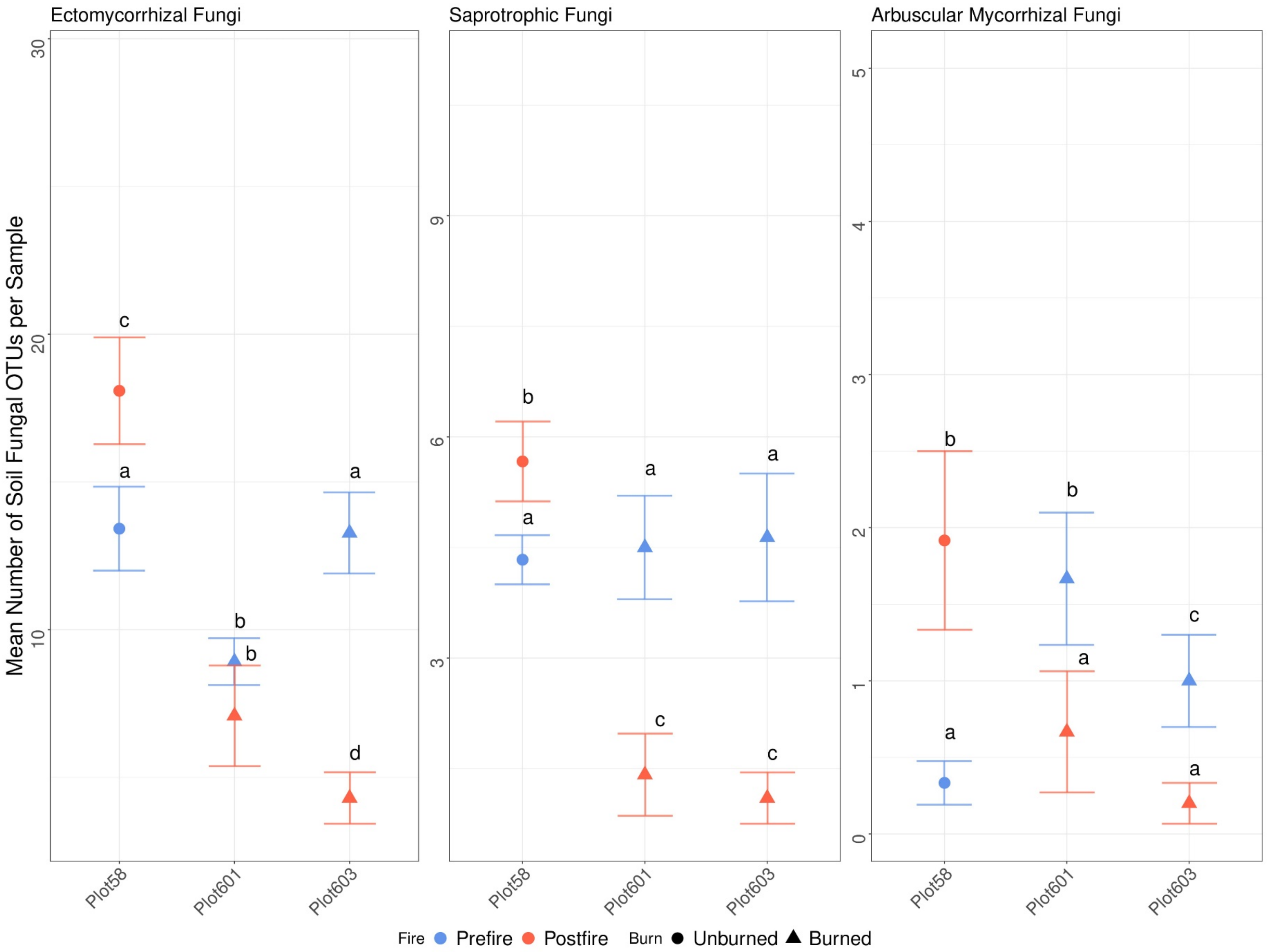

**Figure S6**
NMDS of Bray-Curtis Dissimilarity ordinations comparing (A) bacterial and (B) fungal composition in all plots with colors indicating the 2013 pre-fire and 2016 post-fire samplings and shapes differentiating burned and unburned plots. Ellipses represent 95% confidence interval from the centroid for each group with each group representing a plot. Plot numbers have been placed directly adjacent to their corresponding ellipses.

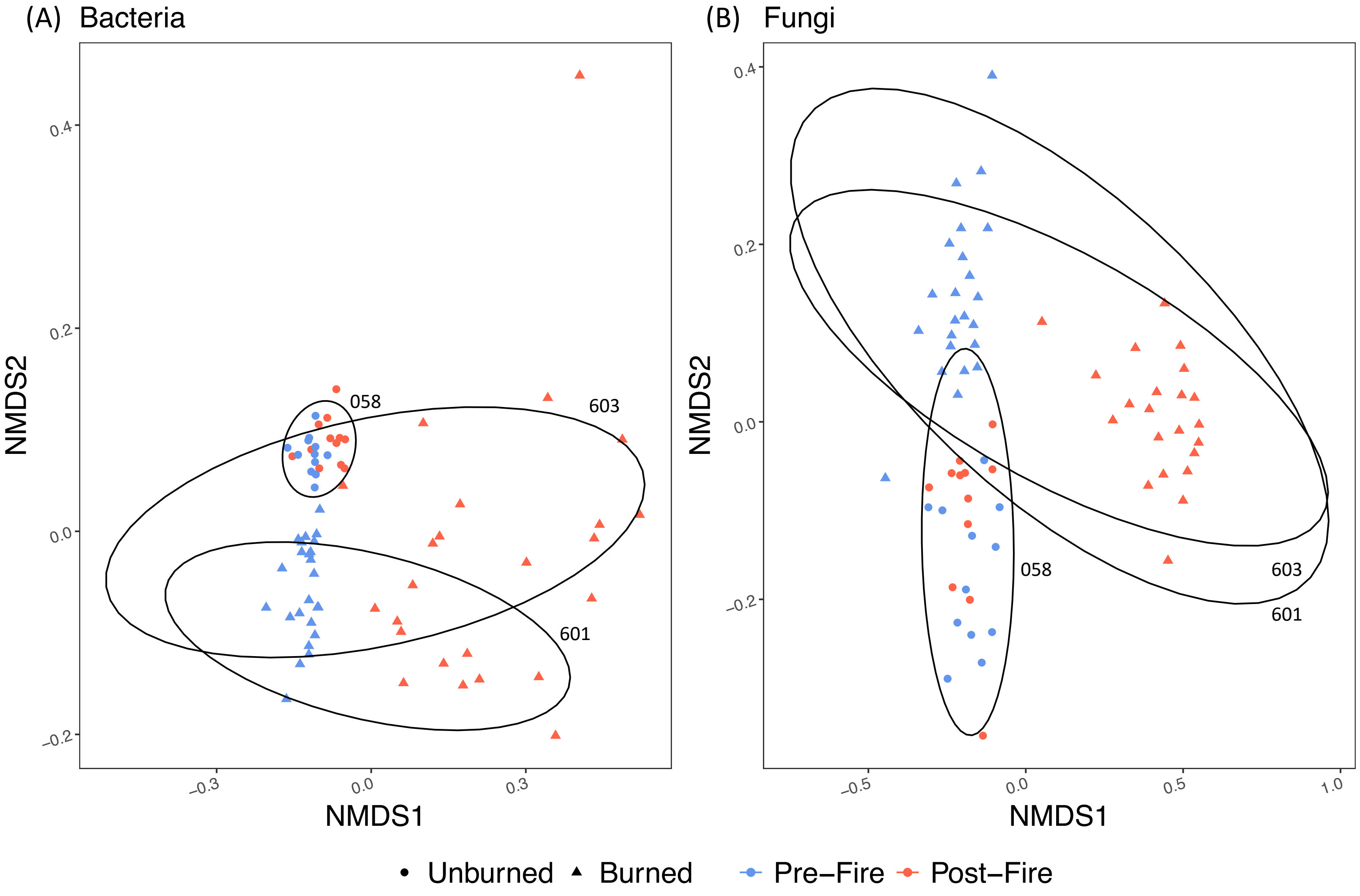
